# An “Aha!” moment precedes the strategic response to a visuomotor rotation

**DOI:** 10.1101/2025.05.11.653302

**Authors:** Max Townsend, Matthew Warburton, Carlo Campagnoli, Mark Mon-Williams, Faisal Mushtaq, J. Ryan Morehead

## Abstract

Strategic behaviour in sensorimotor adaptation tasks is typically described as either a gradual error minimisation process or as a process of learning through trial-and-error. The former predicts a gradual monotonic reduction in error, until some asymptote, while the latter predicts behavioural exploration to discover an efficacious solution. Another explanation is that a sufficiently rich understanding of the task culminates in an “Aha!” moment, and that this richer understanding naturally entails a new strategy. This predicts some period of perseveration in baseline behaviour, followed by an abrupt shift to a new strategic solution. To avoid obfuscation caused by the motor system, we first investigate these hypotheses in a strategy-only aiming game, where participants aim and fire a cannon at targets. Most participants exhibited a period of baseline perseveration followed by a single-trial shift to a distinct strategy. This strategy typically appears to produce the correct compensatory magnitude for a noisily estimated perturbation, but also substantial variability in compensatory sign. We then applied the same analyses to reaching data from a visuomotor rotation task that inhibited implicit adaptation through delaying feedback presentation. Similarly, we found that participants typically perseverated in reaching towards the target before suddenly switching, in a single trial, to good performance in compensatory magnitude. We fit our generative “Aha!” model to 1337 participants across several datasets and show that it faithfully captures these phenomena. Our findings highlight the need to update existing models to account for the abrupt, insight-driven shifts that characterise individual human responses to sensorimotor perturbations.

## Introduction

Correcting for movement errors is integral to the success of an embodied agent. Humans can strategically modify their behaviour to expedite this error-correcting process (Bond & Taylor, 2015; McDougle et al., 2015, 2016; McDougle & Taylor, 2019; Morehead et al., 2011; Taylor et al., 2010, 2014; Taylor & Ivry, 2011, 2012). While recent progress has been made in enumerating core strategic phenomena, and in formulating frameworks from which to investigate these phenomena (Tsay et al., 2024), our understanding of strategic error correction remains shallow.

Traditionally, strategic behaviour in visuomotor rotation (VMR) tasks has been characterised as a gradual error-reduction process. This view predicts a gradual, monotonic increase in learning that tends towards some asymptote, typically modelled by a state-space model (SSM; Coltman et al., 2019; McDougle et al., 2015; Taylor & Ivry, 2011; Takiyama et al., 2015; Zhang et al., 2025). While consistent with group-averaged learning curves, it is unclear if the individual timecourse of learning evolves in the same manner. Indeed, this gradual monotonic group curve can result from averaging together individual learning trajectories that each follow distinct underlying dynamics (Bahrick et al., 1957; Gallistel et al., 2004).

As such, we should also consider that strategic behaviour may instead reflect trial-and-error learning, where actions are selected based on their expected value or inferred efficacy. Here, the agent explores action space (or hypothesis space) to discover an efficacious solution. In this view, the evolution of behaviour is either the result of a reinforcement learning (RL) process, or of updating a set of hypotheses about the context, from which the efficacy of actions is inferred (Collins & Koechlin, 2012; Darshan et al., 2014; Donoso et al., 2014; Griffiths et al., 2010, 2019; Piantadosi et al., 2016; Taylor & Ivry, 2011, 2012; Tsay et al., 2024; Nikooyan & Ahmed, 2015; Izawa & Shadmehr, 2011; Sugiyama et al., 2023; Takiyama et al., 2015). Learning in this manner typically generates clear signals of behavioural exploration before settling on a strategy. These behavioural patterns are inconsistent with, and dissociable from, gradual error reduction.

Importantly, the RL and hypothesis-testing accounts are both incomplete. They assume that the agent’s initial task representation entails a strategy space that is conducive to discovering efficacious strategies. As such, we propose that a key component of strategic learning, ignored by extant theories, is the initial process of changing one’s representation of the task and how one’s goals relate to the task. Moreover, this initial updating of the task representation may naturally entail an efficacious strategy and eat much of the lunch of the trial-and-error accounts.

Indeed, there is much research on the “Aha!” moment. This “aha!” moment may reflect a gain of insight—a qualitatively richer understanding of the task and its demands—that proceeds from reorganising one’s internal representation of the problem (Carpenter, 2020; Jones, 2003; Knoblich et al., 1999, 2001; Kounios & Beeman, 2009; Sprugnoli et al., 2017). This kind of sudden learning has been observed in many areas of cognitive problem solving, such as the compound-word remote associations task (Jung-Beeman et al., 2004), anagrams (Novick & Sherman, 2003), rebus puzzles (Salvi et al., 2016), the nine-dot problem and it’s variations (MacGregor et al., 2001), among others (e.g. Sawyer, 2012; Vartanian & Goel, 2007). Furthermore, we know from work by Manley et al (2014) that awareness of critical task properties may be required in learning to change reach directions, and work on consciousness suggests that this awareness may be all-or-none, rather than gradual (Sergent & Dehaene, 2004).

The strategic problem presented during the baseline (veridical) phase of a VMR task may be simply formulated as the requirement to aim at the target before firing. A change in the action-feedback mapping renders this parsimonious and effective representation uninformative as to how to act successfully. By reorganising this representation to embody more semantic and relational information, an appropriate solution can be brought psychologically closer (Trope & Liberman, 2010). For example, when the action-feedback mapping is modified, a representation of the task that encodes a relationship between the aimed direction and the feedback direction would more naturally lead to an efficacious aiming solution. Learning would manifest, behaviourally, as a period of perseveration of baseline behaviour, followed by a sudden, substantial shift in behaviour following the “Aha!” moment. Indeed, this idea has been raised before in the VMR literature (Taylor & Ivry, 2012; Tsay et al., 2024).

To date, it has remained difficult to elucidate the true nature of strategic behaviour in VMR tasks as extant task designs tend to leave strategic behaviour inextricably linked to implicit adaptation and other contaminants of the motor system. This is either because one measure is used to infer the output of two or more processes, where it is difficult to correctly assign credit to these underlying processes (e.g. Hegele & Heuer, 2010), or because these processes interact in a way which we do not currently understand. To make matters worse, different methods of probing strategic behaviour generate distinct patterns of results (Langsdorf et al., 2021; Maresch et al., 2021a; Maresch et al., 2021b). This is likely because each method changes the nature of the interaction between the strategic system and the motor system. A different approach is to first understand each subprocess in relative isolation, such that we can use the findings as priors when studying the whole process. We can already remove implicit learning from the strategic context by presenting consistent clamped feedback which is irrelevant to performance (Morehead et al., 2017). We aim to complement this by isolating strategic learning from the motor-learning context, while enforcing the same strategic demands.

Here, we investigated aiming behaviour in a series of cannon-shooting tasks where the participant had to overcome a rotation of the visual feedback (Figure 1a). We found that aiming behaviour evolved in a manner that was consistent with an “Aha!” moment, not merely trial-and-error learning or gradual error minimisation. This pattern of baseline perseveration, followed by a delayed single-trial shift to good performance in magnitude is clear both visually and through formal model comparison. We quantitatively confirmed that our task presented similar strategic demands as in a standard VMR task and then carried out the same modelling analyses on a previous dataset from a delayed-feedback VMR task (Brudner et al., 2016) as well as on the comparator study itself (Bond & Taylor., 2015). We found the same pattern of results, where almost all participants were better described by an “Aha!” Moment than by trial-and-error learning or gradual error minimisation.

**Figure 1.**
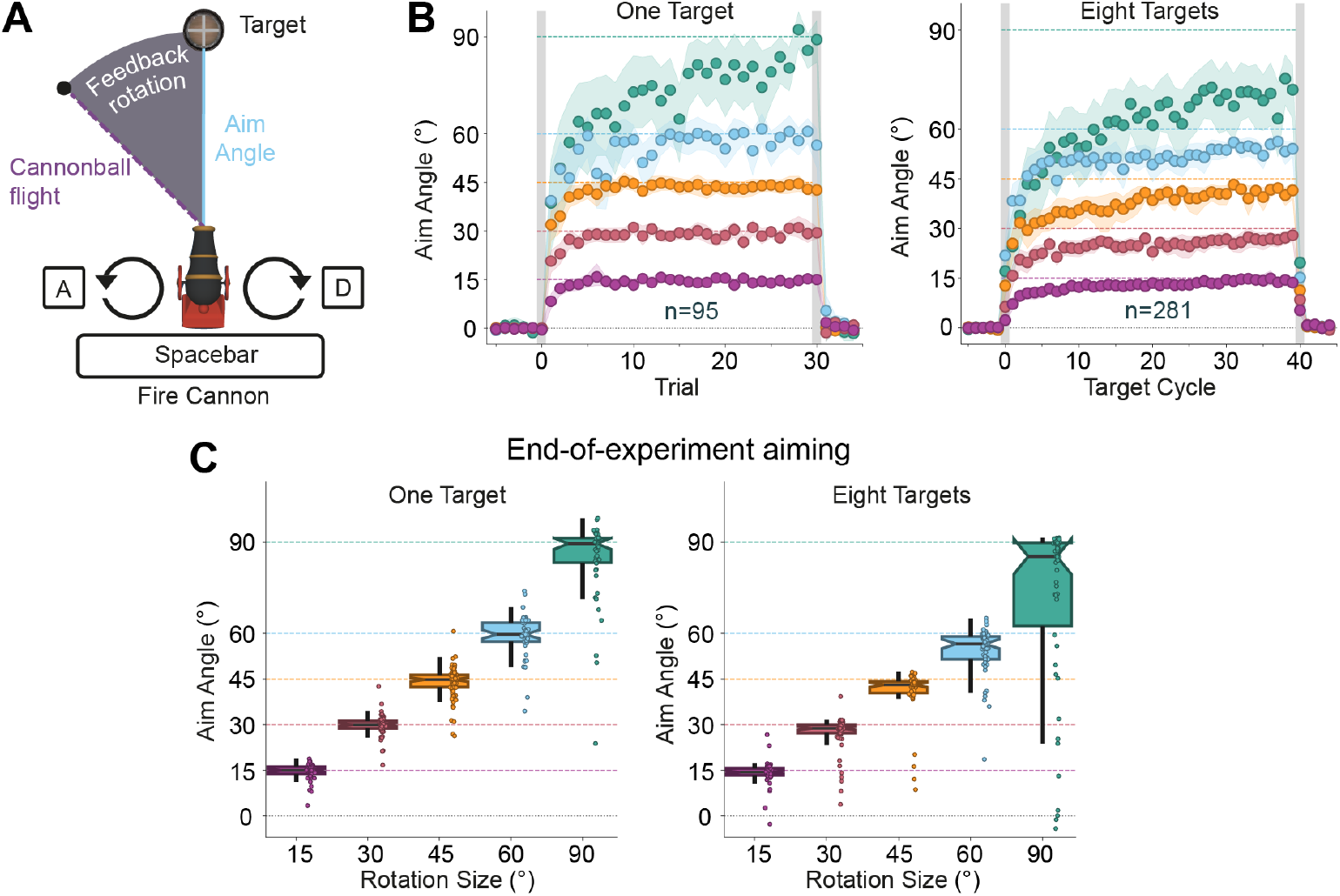
(A) The participant must aim the cannon to shoot a target. The cannon is rotated using the ‘A’ and ‘D’ keys and fired using the ‘Spacebar’ key. (B) Aiming behaviour over the course of experiments 1a, with one target location (left), and 1b, with eight target locations (right). Points show group means, error bands show 95% CIs, and dashed lines indicate ideal aim angles. (C) Aim angles during the final trial or cycle, for experiments 1a (left) and 1b (right), respectively. The boxplots show the quartiles of the data, the whiskers bars show 1.5x the interquartile range, and the points show individual data.

## Methods

### Participants

A total of 761 humans (316 males, 440 females, 5 other genders, age = 34.1 ± 11.8 years) from the United Kingdom and the United States were recruited via the Prolific Academic recruitment platform.

### Inclusion and ethics

Ethical approval was obtained from the University of Leeds school of Psychology Research Ethics Committee (ref PSYC-81). All participants gave informed consent prior to starting the study.

### Experimental Apparatus

Participants aimed a cannon and shot targets using keyboard keys. All participants completed the task remotely on their own desktop or laptop computers, with no further constraints on the hardware they used. The experimental task was constructed using the Unity game engine and default libraries, with custom scripts written in C# and data saved to a remote database.

### Aiming Task

Participants were instructed to rotate the cannon by pressing the ‘A’ and ‘D’ keys, and to fire it with the ‘Spacebar’ key (Figure 1a). They were told to hit as many targets as possible and were not encouraged to act quickly. Tapping the rotation keys rotated the cannon by 1° increments and holding down these keys rotated the cannon at the speed of 125°sec-1. An aiming reticule indicated the aimed location, whose radial distance was held constant (equal to the radial distance of the target). Upon firing, the cannonball flight lasted 300 ms and would terminate at the same radial distance as the target. The target had a radius of 3°, and the cannonball had a radius of 0.5°. Upon being hit, the target exploded and was accompanied by a green ‘Hit!’ text lasting 500 ms; if missed, a ‘buzzing’ sound accompanied red ‘Miss!’ text lasting 500 ms. The Cannon Game task would only play in full-screen mode. At any point, if the participant left full-screen mode, the task was paused and the participant was presented with a pause screen asking them to enter full-screen when they were ready to resume.

Participants were told ‘the cannonball may not always go exactly where you are aiming’. The direction of the shot was subject to noise of similar magnitude to motor noise in reaching tasks (Figure 1b), such that the cannonball endpoint feedback was

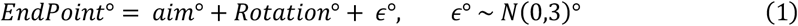

where the integer *aim* is the angle aimed, *ϵ* is gaussian noise, and *Rotation* is the feedback rotation applied to the endpoint feedback. At the start of each trial, the cannon direction was sampled from a uniform random distribution spanning 60° either side of, but at least 5° away from, the angle equal to − *Rotation*° (i.e., the ideal aim angle).

#### Experiment 1

The first experiment sought to explore how participants updated their aiming behaviour in a simple task which requires the participant to aim at targets, much like in a visuomotor rotation (VMR) task.

In experiment 1a (n = 98), the target was presented in the same position (12 o’clock) on every trial. The experiment comprised six blocks, where each block included 30 rotation trials with a rotation applied to the direction of feedback. The rotation phase was sandwiched between 15 baseline and 15 washout trials, each with veridical feedback (Figure 1a). Within each block, the rotation phase presented a single feedback rotation for all trials, such that one group of participants each experienced every sign/magnitude combination of {±15, ±30, ±45}°, and the other group experienced all of {±45, ±60, ±90}° by the end of the six experimental blocks. Three participants were excluded for having >20% trials with <100ms aiming time.

In experiment 1b (n = 301), the target was presented in one of eight locations, evenly spaced in 45° increments from the target at 12 o’clock. The order of target location was pseudorandomised, where each location occurred once in each eight-trial cycle. There was a single block of 40 cycles of rotated feedback, sandwiched between five baseline and five washout cycles. During the rotation phase, each participant experienced one rotation, sampled from {±15, ±30, ±45, ±60, ±90}°. Twenty participants were excluded for having >20% trials with <100ms aiming time.

#### Experiment 2

Experiment 2 implemented aiming requirements which were much more like those in VMR tasks. To this end, it included an autocorrection module (i.e., a spatially-generalising SSM adapted from McDougle et al. (2017)) which learns a bias that is added to the feedback direction. This bias gradually minimises the difference between the aimed location and the endpoint feedback (Figure 2a & b). The resultant formula for the endpoint feedback direction is

**Figure 2.**
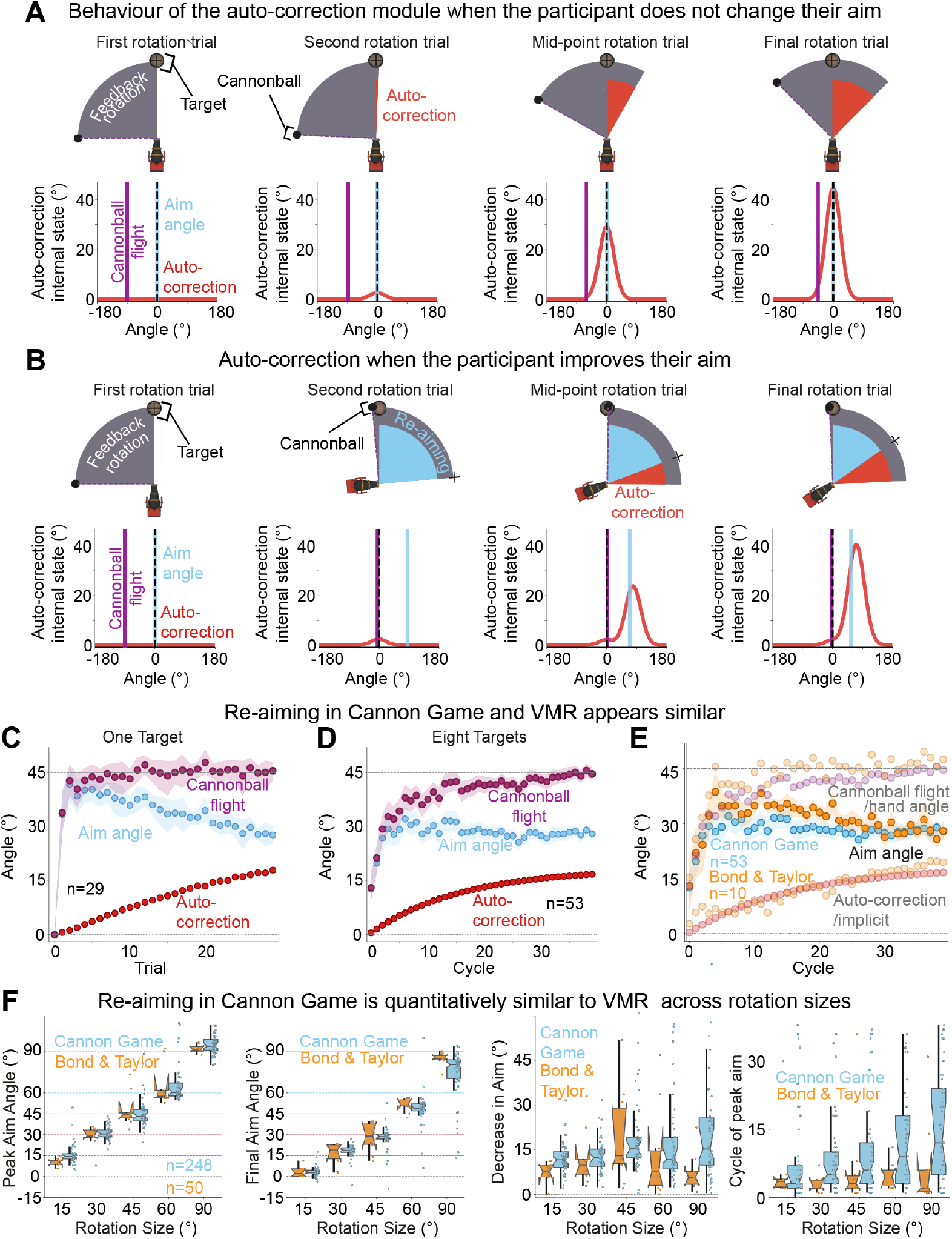
In experiments 2a and 2b, the cannon gradually learned to autocorrect for the feedback rotation. (A) & (B) The direction of cannonball flight (purple) is equal to autocorrection (red) + re-aiming (blue) - feedback rotation (grey). Timecourse of learning for the auto-correction module when a participant doesn’t adjust their aim, and timecourse of learning when a participate learns to re-aim, respectively. Top row: The contribution of each learning system (human re-aiming and cannon autocorrection), and the task outcome, on a given trial. Bottom row: internal state of the auto-correction module (red) generalises around the workspace. Performance in the 45° condition, under a one-target (C) or eight-target (D) experiments. (E) Performance during the rotation phase in experiment 2b (blue) compared to data from Bond & Taylor (2015; orange). Auto-correction model (SSM) parameters were fit to implicit learning in Bond & Taylor; see methods section. Autocorrection/implicit learning and cannonball flight/hand angle are included for transparency but de-emphasised as the aiming comparison is primary. (F) Comparison of summary statistics for each rotation size. In order, from left to right: each participant’s greatest cycle-averaged aiming value; each participant’s aiming value on the final rotation cycle; each participant’s final aim subtracted from their peak aim; the cycle number at which each participant’s peak aim was observed. For (C-E), points show group means and error bands show 95% confidence intervals. For (F), the boxplots show the quartiles of the data, the whiskers bars show 1.5x the interquartile range, and the points show individual data.

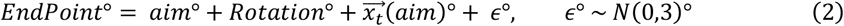

where 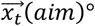 is the output of the autocorrection module for integer angle *aim* on trial t.

The state vector 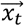 is updated after each trial t according to

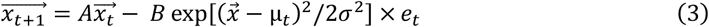

where 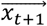 is the state vector on trial t+1, *A* is the retention factor, *B* is the learning rate, *e*_*t*_ is the prediction error (*EndPoint*_*t*_ − *AimAngle*_*t*_), 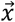 is the ordered vector of all integer angles around the workspace (−179,−178…,180), and μ_*t*_ is the centre of gaussian generalisation on trial t. The parameter μ_*t*_ is set to the aim angle on trial t, such that the generalisation of learning is centred on the aimed angle (Figure 2a & b). Parameter *σ*, which defines the width of generalisation, was set to 30° (selected to be consistent with previous generalisation widths (Krakauer et al., 2000; Mattar & Ostry, 2007; McDougle et al., 2017; Taylor & Ivry, 2013)).

For Experiment 2a (n = 61), the parameters *A* and *B* were set to 0.99 and 0.0275 respectively (consistent with the values found in McDougle et al. (2015)). The target was presented in a single location and the task comprised six 60-trial blocks, each including a 30-trial rotation phase sandwiched between a 15-trial baseline phase and 15-trial washout phase. One group experienced one of {±15, ±30, ±45}° in each block, and the other group experienced one of {±45, ±60, ±90}°, akin to experiment 1a. One participant was excluded for having a mean absolute aim of <3° (i.e., the target radius).

For Experiment 2b (n = 301) we wanted to approximate actual aiming requirements experienced by participants of a VMR study. To do this, we fit the learning rate (*A*) and retention rate (*B*) parameters of the autocorrection module to data from Bond & Taylor’s (2015) eight-target reaching task. Parameter fitting was carried out by finding the maximum likelihood over all participants within each rotation size separately, using the optimize.minimize function (with the ‘L-BFGS-B’ method) of the scipy (scientific python) package for Python (parameter fits in Table 1).

**Table 1.** Learning rate and retention rate parameter values for the autocorrection module used in experiment 2b. These are the result of fitting to data from Bond & Taylor (2015).

| Rotation<br>($^\circ$) | Autocorrection<br>module learning | Autocorrection<br>module retention |
| --- | --- | --- |
| 15 | 0.0385 | 0.9985 |
| 30 | 0.0152 | 0.9956 |
| 45 | 0.0152 | 0.9951 |
| 60 | 0.0118 | 0.9875 |
| 90 | 0.0044 | 0.9918 |

As in experiment 1b, the target was presented in one of eight locations, evenly spaced in 45° increments around the workspace. The order of target location was pseudorandomised, where each location occurred once in each eight-trial cycle. There was a single block of 40 cycles of rotated feedback, sandwiched between five baseline and five washout cycles. During the rotation phase, each participant experienced one rotation, sampled from {±15, ±30, ±45, ±60, ±90}°. Twenty-seven participants were excluded for having >20% trials with <100ms aiming time, and three participants were excluded for having a mean absolute aim of <3° (i.e., the target radius).

### Data Analysis

Analyses were completed with custom scripts written in Python 3.7. We used an α-value of 0.05, such that any p-value less than α indicates a statistically significant difference. Unless stated otherwise, all t-tests were two-tailed and Bonferroni corrections were applied for multiple comparisons. Our primary dependent variable was the signed difference between the aim angle and the target angle, with the resulting angle multiplied by −1 for positively signed rotations so that a positive angle was one that counteracted the applied perturbation. Fifty-four participants were excluded (all previously listed n values were calculated before participant exclusion) because they met at least one of the following criteria: fifty participants were excluded for having >20% trials with <100ms aiming time, and four participants were excluded for having a mean absolute aim of <3° (i.e., the target radius). There was no trial-level exclusion

In experiments 1 & 2, we used t-tests to check that baseline aiming was not significantly different from zero, and to check whether aiming was significantly different from zero during mid learning and late learning. We also tested whether the washout aiming (excluding the first trial/cycle) significantly differed from zero. In experiments 1a & 1b, t-tests were used to assess whether aiming was significantly different from the ideal aim angle during mid and late learning. In experiments 2a & 2b, we instead tested if the ‘final cannonball angle’ (aim + offset learned by autocorrection module) was significantly different from the ideal cannonball angle during late learning and mid learning. We also applied a t-test on the implicit autocorrection vs zero during washout. In experiments 1a & 2a, mid learning was defined as the middle five rotation trials, and late learning was defined as the final five rotation trials. In experiments 1b & 2b, mid learning was defined as the middle five rotation cycles, and late learning was defined as the final five rotation cycles. In experiments 1a and 2a, where participants encountered a given rotation magnitude twice with opposite signs, data was averaged over both blocks.

For the direct comparison to VMR data, we compared data from experiment 2b to data from Bond & Taylor (2015). A 2 (experiment type) x 5 (rotation size) x 3 (learning phase) ANOVA was carried out, where experiment type was denoted whether the experiment was ours or from Bond & Taylor, and learning phase was either early learning (first 10 rotation cycles), mid learning (middle 10 rotation cycles), or late learning (final 10 rotation cycles). We also applied t-tests to compare the following properties across experiments: the cycle of peak aiming, the magnitude of peak aiming, and the decrease in aiming from peak to final rotation cycle.

We then go on to investigate behaviour that has been realigned to each individual’s “Aha!” moment. We use paired t-tests to see if the pre-”Aha!” rotation trial has aiming that is significantly different from baseline, and we also use one-sample t-tests to test If the “Aha!” trial aiming magnitude is different from the ideal aiming magnitude. We also used t-tests to detect any significant change in trial-to-trial variability between the baseline phase and the pre-”Aha!” rotation phase, or if there was any significant error correction in the pre-”Aha!” rotation phase.

### Data-driven “Aha!” moment detection

To detect the “Aha!” moment, defined as the first changepoint in aiming behaviour during the rotation phase, we applied a non-parametric, model-free changepoint algorithm to each individual’s aiming magnitude trial series. We used the Pruned Exact Linear Time (PELT; Killick et al., 2012) implemented in the rupture (version 1.1.7) Python library (Truong et al., 2020). The PELT algorithm minimises a penalised cost function over possible segmentations. The cost function employed a radial basis function (RBF) kernel to measure dissimilarity between proposed segments, handling arbitrary shifts and the potentially multimodal nature of behaviour after the “Aha!” moment. For all datasets in the main analyses, we used the same parameter settings: the penalty (‘pen’), which determines the penalty of each detected changepoint (higher values lead to fewer detected changepoints), was set to 0.8; the minimum segment size (‘min_size’) was set to 1 (such there need not be multiple trials between each changepoint); the ‘jump’ parameter was set to 1, to ensure that a segmentation was considered at every trial.

The first changepoint during the rotation phase was then selected as the “Aha!” trial. In the supplementary analysis of Ding et al. (2025; Figure S4), ‘pen’ was set to 0.2 for greater sensitivity to changes, but all other parameters were left unchanged. Importantly, during the rotation phase, participants in Ding et al. were explicitly instructed that cursor feedback would no longer correspond to their movements and that they should discover the movement that correctly results in the cursor being placed on the target. This instruction appears to have expedited the “Aha!” moment for many participants; as such, for some analyses (e.g., Figure S4B & F) we exclude 145/560 participants who pre-emptively changed their aim angle on the first rotation trial (i.e., before observing any rotated feedback).

### Computational models

As part of our investigation into the individual timecourse of aiming, we compare three models of noisy aiming: a single-process SSM to model monotonic, gradual learning; a bespoke Q-Learning model of trial-and-error learning; and the Q-Learning model embedded in a two-state HMM as the “Aha!” moment model.

### State-space model

The state of the gradual-learning model, *x*_*t*_, determines the aim on trial *t*. We assume that the process of executing an intended plan is inherently noisy, such that the aim on trial *t* is defined

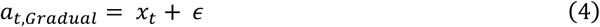

where *a*_*t,Gradual*_ is the aim on trial *t, x*_*t*_ is the internal state of the SSM, and the execution noise *ϵ* follows an unbiased normal distribution

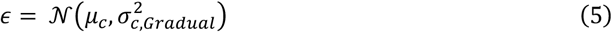

defined by *μ*_*c*_ = 0, and execution variance 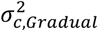. The state is updated according to

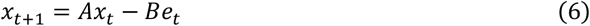

where *A* is the retention rate, *B* is the learning rate, and *e*_*t*_ is the prediction error on trial *t*, as defined in equation 3. The SSM has three free parameters: *A, B*, and 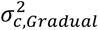. We also tested a dual-state SSM, but found that that produced worse complexity-penalised fits than the single-state SSM defined above (Figure S3).

### Trial-and-error learning model

To represent pure trial-and-error learning, we implemented a Q-learning model which learns from dense (error-based) rewards and learns sign and magnitude separately via off-policy updates. The model assumes each trial is independent, with no state transition function, and so has a one-step horizon. This statelessness is justified because feedback is immediate on each trial, and the action-reward mapping is independent of action history. The action policy is updated to minimise expected feedback errors, where the action space has been hierarchically discretised into magnitudes of 1° increments *M* = {*a*_0_, …, *a*_180_} and sign *SGN* = {+, −}. The model maintains separate value functions across trials: *Q*_*mag*_: *M* → ℝ and *Q*_*sgn*_: *SGN* → ℝ (both initialised to zero). The policy *π* is hierarchically defined via two softmax functions:

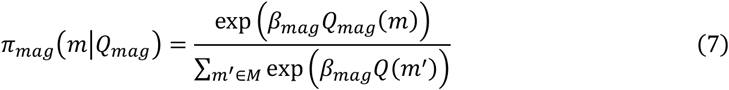

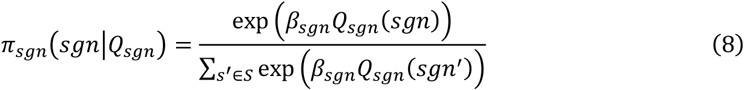

where *m* is the candidate magnitude, *sgn* is the candidate sign, *β*_*mag*_ is the inverse temperature for magnitude, and *β*_*sgn*_ is the inverse temperature for sign.

#### Action Selection

The model generates actions by sampling from the hierarchical mixture policy over the expanded action space:

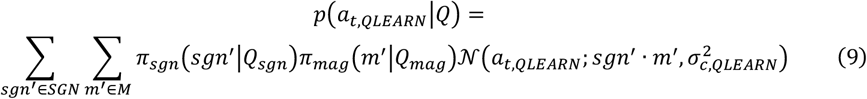

where *a*_*t,QLEARN*_ is the aim on trial *t* and 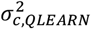 is the execution variance.

#### Update Step

After observing feedback, the model performs off-policy Q updates using the temporal difference (TD) error. This off-policy update is analogous to counterfactual reasoning under the assumption that the action-feedback mapping is uniform across the workspace (e.g., “what would the outcome have been if I had aimed there instead?” and “where would have been the best direction to have aimed?”). Each update utilises the maximum attainable reward for the specific sign or magnitude being updated, where reward is defined

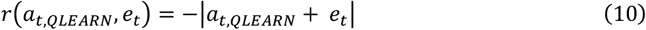

where the prediction error *e*_*t*_ is counterfactually assumed to be the same for all aim angles. For sign updates, compute the maximum reward attainable (*maxR*_*sgn*_) using each sign *s* ∈ *S*:

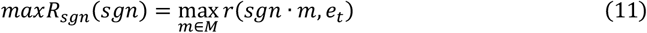

then compute the TD error *δ*_*sgn*_ and update *Q*_*sgn*_:

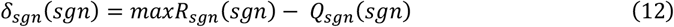

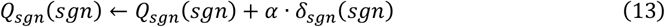

where *α* ∈ (0,1] is the learning rate. A similar process is then applied to update *Q*_*mag*_; for each magnitude *m* ∈ *M*, compute the maximum reward availalable *maxR*_*mag*_ over either sign and update with the TD error *δ*_*mag*_:

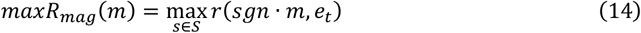

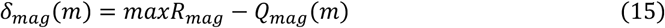

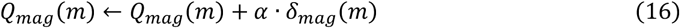

This Q-learning model has four free parameters: 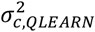, *α, β*_*mag*_, and *β*_*sgn*_.

### The “Aha!” model

We hypothesise that humans first require an “Aha!” moment to realise that a new aiming solution must be generated. To implement this, we extend the Q-learning model to be part of a two-state Hidden Markov Model (HMM), where the learner transitions between a veridical-feedback state *s*_*v*_, where no learning occurs, and a perturbed-feedback state *s*_*p*_, where learning is governed by the Q-learning model defined above. The HMM enables a potentially abrupt transition from *s*_*v*_ to *s*_*p*_ (and back). This transition may be arbitrarily delayed, as determined by the initial state probabilities **P**_**0**_ = [1,0] and symmetrical prior transition matrix defined:

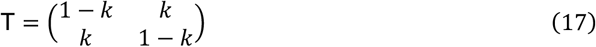

where *k* ∈ [0,1) is the prior transition probability (low values favour persistence in the current state). The prior predicted state distribution at trial *t* is therefore **P**_***t***|***t***−**1**_ = **P**_***t***−**1**_T.

#### Action Selection

This Q-HMM “Aha!” model first samples the hidden state *S*_*t*_*∼***P**_*t*|*t*−1_, where *S*_*t*_ ∈ {*s*_*rel*_, *s*_*irr*_}, then an intended aim, conditionally on *S*_*t*_. Specifically, for *S*_*t*_ = *s*_*irr*_, the sensory prediction error is considered irrelevant to the agent’s goals, so the planned aim is 0°. For *S*_*t*_ = *s*_*rel*_, where the prediction error is assumed relevant to the agent’s goals, the aim is sampled from the hierarchical mixture policy *a ∼ π*(· |*Q*_*mag*_, *Q*_*sgn*_) where *π*(*a*|*Q*_*mag*_, *Q*_*sgn*_) = *π*_*mag*_(|*a*||*Q*_*mag*_) · *π*_*sgn*_(*sgn*(*a*)|*Q*_*sgn*_) using equations 7 and 8. Thus, on trial *t*, the action *a*_*t*,AHA_ is drawn from the state-marginalised predictive distribution defined by:

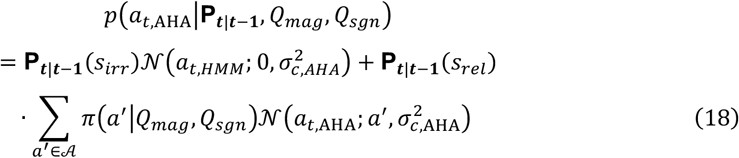

where 𝒜 = {−180, …, 179} uses the same discretised action space as the base Q-learning model, and 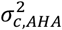 is execution variance.

#### State Inference and Update Step

The sensory prediction error (*e*_*t*_) serves as the context emission that drives the posterior state distribution **P**_***t***_. The state-conditioned emission probability *p*(*e*_*t*_|*s*_*t*_) evaluates how well the predicted aim distribution of each state would minimise the prediction error under contextual uncertainty 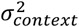 (i.e., uncertainty over the prediction error):

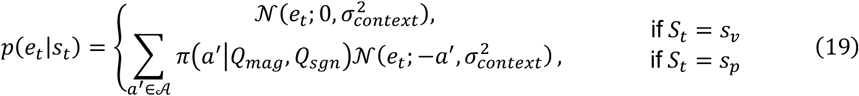

The posterior is then given by

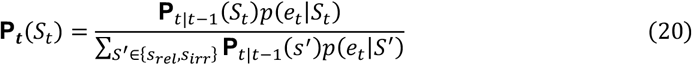

for all *S*_*t*_ ∈ {*s*_*rel*_, *s*_*irr*_}. The Q functions (*Q*_*mag*_, *Q*_*sgn*_) are updated using the off-policy Q-learning rule as in eq. x, but conditioned on the posterior probability of being in the prediction-error-is-relevant state **P**_***t***_(*S*_*t*_ = *s*_*rel*_). This conditioning means that learning may stay effectively at zero until an abrupt “probability explosion” occurs (often within a single trial; Figure S1). This “probability explosion” is triggered when the posterior probability of being in state *s*_*rel*_ jumps from very small (i.e., <<1e-5) on trial *t-1* to ≈ 1 on trial *t* (i.e., from **P**_***t***_(*S*_*t*_ = *s*_*rel*_) ≈ 0 on trial *t-1* to **P**_***t***_(*S*_*t*_ = *s*_*rel*_) ≈ 1 on trial *t*). This means that the Q-functions (*Q*_*mag*_, *Q*_*sgn*_) on trial *t* receive a full update, allowing one or both dimensions of the policy on trial *t*+*1* to to be centred on the ideal value. This is akin to a human experiencing an “Aha!” moment, after observing feedback on trial *t*, and then implementing their new strategy on trial *t*+*1*. See Figure S4A for actual human trial-series aiming data overlaid on model simulations using fitted parameters.

The model is not intended to be an exhaustive mechanistic account; instead, it abstracts away some of the more ethereal components that lead up to the “Aha!” moment by overloading the state inference architecture. Put simply, the state inference architecture is perhaps a partial black-box proxy for the poorly understood processes that culminate in an “Aha!” moment.

The “Aha!” model has six free parameters: 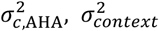, *α, β*_*mag*_, *β*_*sgn*_, and *k*.

### Implicit learning extension

To apply these models in tasks with implicit learning, we extend all three with the same implicit learning component defined in equation 3. In this way, the implicitly learned bias on trial *t*, 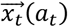, is added to the feedback rotation, such that the feedback on a given trial is computed:

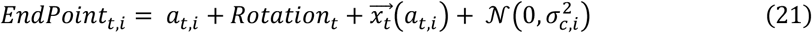

where *i* ∈ {*SSM, QLEARN*, AHA} is the model being used. Each model then rolls out as previously defined with this modular implicit learning added to the feedback. This implicit module does introduce a direct dependency on past behaviour, but it is negligible, so we keep the stateless nature of the Q-learning model (and its instantiation within the “Aha!” model). There are now three additional free parameters during the fitting process of each model to the Bond & Taylor dataset; the implicit learning rate *A*_*imp*_, the implicit retention factor *B*_*imp*_, and the standard deviation of implicit gaussian generalisation *σ*_*imp*_. When fitting to the 8-target Cannon Game-with-autocorrection dataset, these implicit parameters were fixed to the values used by the autocorrection module in the task (Table 1).

All models were fit to the first block of each participant separately, for each experiment separately. Subsequent blocks were excluded as behaviour there may be contaminated by experience from previous blocks. Parameters were optimised through maximum likelihood estimation using the L-BFGS-B algorithm (from the scipy.optimize package). Each model was fit 500 times to each participant, each time using pseudo-random initial parameter values generated through latin hypercube sampling over the entire parameter space.

## Results

### Experiment 1

We set out to characterise aiming behaviour in a shooting task, where participants must aim and fire a cannon to hit targets (Figure 1a). This experiment was designed to explore aiming in its purest form; isolated from any implicit motor-learning system (such as adaptation). In condition a of this experiment, participants were exposed to either the ±15°, ±30°, and ±45° rotations, or the ±45°, ±60°, and ±90° rotations, with only one target location. Participants quickly learnt to fully compensate for all rotation sizes by aiming in the opposite direction (Figure 1b, left).

During the baseline phase, prior to all rotations, there was no bias in aiming (t-tests against zero; Bonferroni-corrected ps > 0.334). By the final five rotation trials of the block for all rotation sizes, participants had learned to respond to the feedback rotation by aiming in the opposite direction (Figure 1c, left; t-tests against zero; ps < 0.001). By the final five rotation trials, the 90° group still aimed differently to the ideal aim angle (95%CI, 80.12°, 88.33, t(45) = 2.83, *p* = 0.034), but aiming in all other rotation groups was not different from the ideal aim angle (t-tests against ideal aim angle; ps > 0.146). For all rotations (ps > 0.467), except 90° (95% CI, 62.75°, 81.98°, t(45) = 3.69, p = 0.002), this ideal aim angle was already achieved by the middle five trials (trials 13 to 17). After the first trial of the washout phase, there was no residual bias in aiming (t-test against zero; ps > 0.103).

In condition b, five new groups of participants were each exposed to one of ±15°, ±30°, ±45°, ±60°, or ±90° rotations. With eight target locations, this condition comprised a single block presenting one rotation size. For all rotation sizes, participants learnt to fully compensate for the rotation. For all groups, there was no bias in baseline aiming (ps = 1), and by the final five cycles of the rotation phase, aims had significantly changed from zero (Figure 1c; ps < 0.001). During this late-rotation phase, aiming was smaller than the ideal aim angle in the 30° group (95% CI, 25.4°, 28.65°, t(57) = 3.67, p = 0.002), the 45° group (95% CI, 38.95°, 43.09°, t(56) = 3.86, p = 0.001), the 60° group (95% CI, 52.17°, 56.36°, t(56), p < 0.001) and the 90° group (95% CI, 61.7°, 88.32°, t(53) = 5.48, p < 0.001), but not the 15° group (*p* = 0.646). During the middle five cycles (cycles 18 to 22), aiming in all groups was different from the ideal angle (ps < 0.002). After the first washout cycle, there remained a small bias in aiming towards the previously ideal aim angle in the 15° (95% CI, 0.24°, 0.98°, t(54) = 3.33, p = 0.007) and 60° (95% CI, 0.12°, 0.54°, t(56) = 3.1, p = 0.015) groups, but no other (ps > 0.094).

### Experiment 2

The second experiment sought to bring the task requirements closer to those in a VMR task. This was achieved by adding in an autocorrection module to the cannon (Figure 2a & b). The autocorrection module automatically and covertly corrected for task error in a monotonic fashion (akin to the typical SSM formulation of implicit adaptation; see Methods section). Participants learned to fully compensate for all feedback rotations. The initial rapid increase in aiming was followed by a slow decrease in the amount re-aimed, such that the cannonball would persist in going towards the target region as the autocorrection module learns to compensate more (Figure 2c & d).

In condition a of experiment 2, there was one target position, and participants were exposed to either the ±15°, ±30°, and ±45°, or the ±45°, ±60°, and ±90° rotations. During the baseline phase, there was no bias in aiming for any rotation size (ps > 0.441). When combined with the output of the autocorrection module, the resulting cannonball angle during the final five trials was not different from the ideal angle in any rotation group (ps = 1). After the first trial of the washout period, aiming in all rotations was biased in the direction opposite the previously ideal angle (ps < 0.001). This effect was because of the persisting effects of the autocorrection module, whose learned bias now offset the cannonball angle in the opposing direction (ps < 0.001).

In condition b, there were eight target positions, and five new groups of participants were each exposed to one of ±15°, ±30°, ±45°, ±60°, or ±90° rotations in a task comprising a single block. During the baseline phase, there was no aiming bias for any group (ps > 0.356). In the middle five cycles of the task, a t-test versus the ideal angle revealed that participants (combined with autocorrection) already fully compensated for the 15° rotation (95% CI, 14.07°, 15.26°, t(57) = 1.122, p = 1), but not in any other group (ps < 0.02). There was much variability in the cycle index at which participants exhibited their greatest cycle-averaged aim (Figure 2f, rightmost panel); the mean (±standard deviation) number of cycles until peak aim were 8.24 (±10.94), 8.12 (±8.56), 9.98 (±10.7), 12.65 (±10.57), and 15.69 (±11.97) for the 15°, 30°, 45°, 60°, and 90°, respectively, such that people tended to reach their peak aims later in the experiment under larger rotations. During their cycle of peak aim, participants aimed 15.54° (±6.85°), 30.95° (±4.39°), 46.18° (±12.14°), 65.94° (±14.3°), and 94.92° (±7.39°), in their respective groups. By the final rotation cycle, aiming reduced to 3.24° (±3.43°), 18.18° (±4.58°), 27.94° (±8.73°), 47.95° (±11.91°), and 73.68° (±18.41°), respectively.

### Direct comparison with VMR task

To explore whether aiming behaviour is consistent with that observed in VMR tasks, we now directly compare key properties of learning between experiment 2b and the eight-target experiment of Bond & Taylor (Figure 2e & f). We find a general, but not complete, similarity between the two sets of aiming data, indicating that insights discovered in Cannon Game may also apply to the strategic decision-making processes employed in a VMR task.

We first conducted a 2 (experimental paradigm; Explicit vs Reaching) x 5 (rotation size) x 3 (learning phase) ANOVA on aim angles and found a main effect of experimental paradigm [F(1, 933) = 20.111, p < 0.001, η_p_^2^ = 0.021 95% CI [0.006, 0.044]] which varied depending on rotation size F(4, 933) = 8.434, p < 0.001, η_p_^2^ = 0.035 95% CI [0.017, 0.065]]. We found no interaction effect of learning phase x paradigm [F(2, 933) = 1.165, p = 0.312, η_p_^2^ < 0.001 95% CI [0.000, 0.013]] or the three-way interaction [F(8, 933) = 0.745, p = 0.651, η_p_^2^ = 0.006 95% CI [0.004, 0.031]]. Post-hoc pairwise comparisons (simple effect of paradigm at each rotation size) revealed a significant difference of paradigm for the 90° groups (95%CI, 5.8°, 14.7°, t_(23.1)_ = 4.76, *p* < 0.001, Cohen’s d = 1.116), but for no other rotation size (ps > 0.433). Between the two experiments, there was no difference in the cycle index of peak aim for the for any rotation size (ps > 0.129; Figure 2f, rightmost panel). Within this cycle of peak aiming, there was no difference in aiming for any rotation size (ps > 0.085). In the final rotation cycle, there was no difference in aiming between experiments for any rotation (ps > 0.363; Figure 2f, second panel). There was a greater decrease in aiming (the difference between peak aim and final-cycle aim; Figure 2f; third panel) in Cannon Game (12.3°) than Bond & Taylor (6.89°) for the 15° rotation (95% CI, 1.86°, 8.95°, t(66) = 3.05, p = 0.017, Cohen’s d = 1.04), but no other (ps > 0.087).

### Elucidating the “Aha!” moment

Learning at the group level, thus far, appears to evolve in a gradual, monotonic fashion until the ideal aim angle is reached. However, this gradual, monotonic group curve could result from averaging over variably parameterised individual functions which may not be gradual and monotonic themselves (Bahrick et al., 1957; Gallistel et al., 2004). To investigate this, we began with data-driven analyses, which we then followed up with various model-based analyses. In these analyses, we included data from all our experiments, as well as reach-based datasets from Bond & Taylor (2015) and Brudner et al. (2016) to investigate whether our findings generalise to the VMR context (also see Figure S4 for similar analyses on the recent delayed-feedback dataset from Ding et al., 2025).

We began by finding the hypothesised “Aha!” moment for each participant of every dataset via the data-driven PELT algorithm (see Methods section for how exactly this is achieved). For example, in Figure 3A, we can see an exemplar from experiment 1b (left), from Bond & Taylor (middle), and from Brudner & Taylor (right). On these plots, the “Aha!” trial (the first changepoint in the rotation phase; cyan line) clearly demarcates a discontinuity between baseline-like aiming behaviour and some newly adopted aiming strategy. Each exemplar exhibits sign-flip errors in the early section of their post-”Aha!” aiming, indicating that the “Aha!” moment did not result in an efficacious strategy *en toto*.

**Figure 3.**
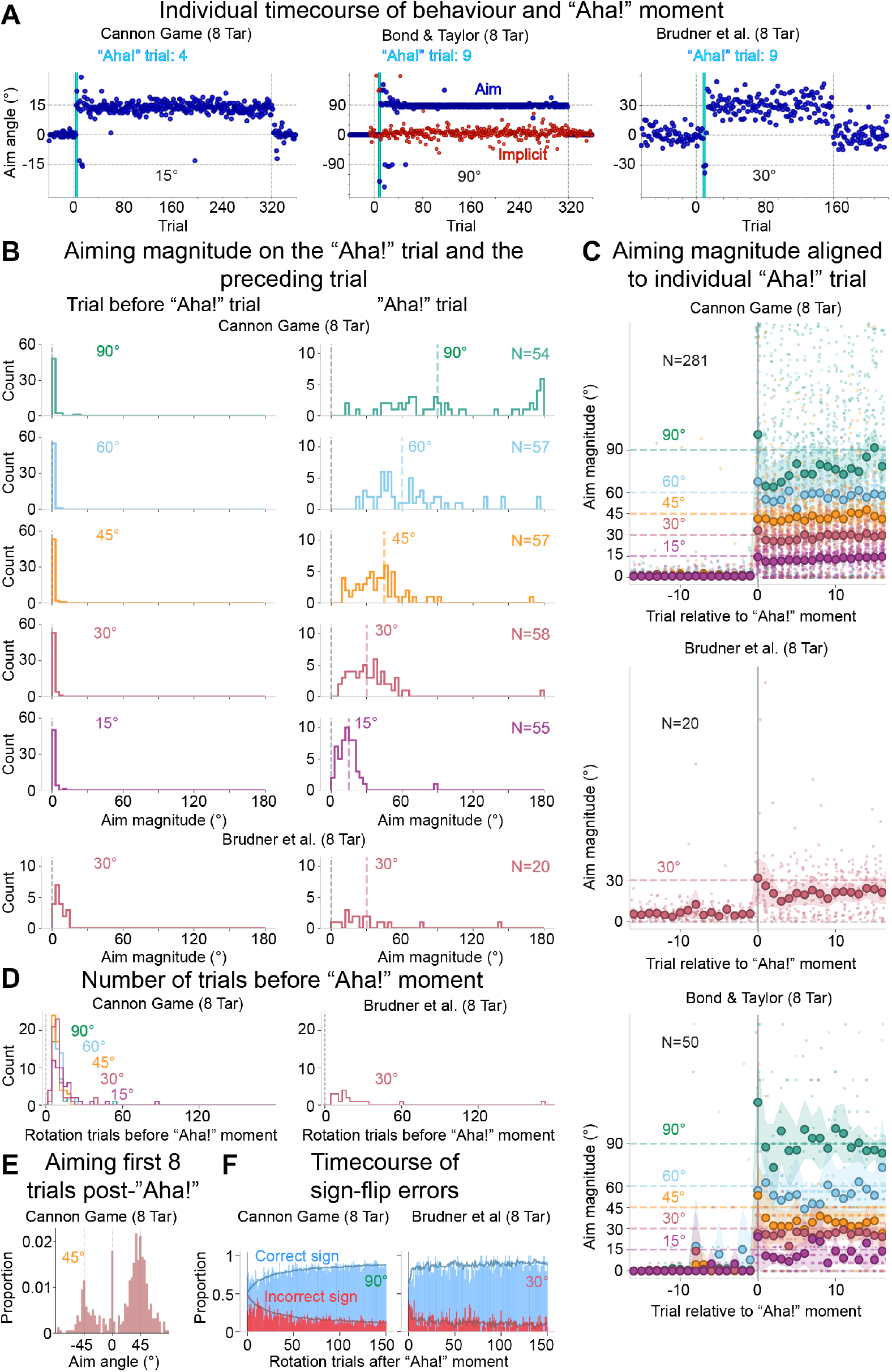
An “Aha!” moment precedes the generation of a new strategy. (A) Examples of individual aiming behaviour (blue) over time in vanilla 8-target Cannon Game (left), Bond & Taylor (2015; middle), and Brudner et al (right). Note the change in y-scale. The algorithmically detected “Aha!” trial is highlighted in cyan (see methods for details on PELT algorithm), and for Bond & Taylor, implicit learning is in red. (B) All participant aim magnitudes for the trial immediately preceding their respective “Aha!” moment (left) and on the subsequent trial (right), for vanilla Cannon Game (top) and Brudner et al (bottom). (C) De-signed aiming magnitude over time, for vanilla Cannon Game (top), Brudner et al (middle), and Bond & Taylor (bottom), where each individual’s data have been realigned such that trial 0 is the first trial at which a new strategy has been observed for that participant (D) Histogram of the number of rotation-phase trials before each participant’s respective “Aha!” moment for vanilla Cannon Game (left) and Brudner et al (right). (E) Distribution of all participant’s aim signed aim angles for the first eight trials following their respective “Aha!” moment (including the “Aha!” trial), for the 45° group of the eight-target vanilla Cannon Game. (F) Proportion of aims having the correct (blue) and incorrect (red) sign over time, during the rotation phase, for vanilla Cannon Game (left) and Brudner et al (right). Solid lines are the “Aha!” model aiming sign proportions. This data is realigned to each individual’s respective “Aha!” moment and includes only the aims which were >15deg away from the target. Error bands = 95% confidence intervals.

To reduce the dimensionality of the problem, and to address the fact that the sign and magnitude of the perturbation may be learned separately, we de-signed the aiming data, allowing the investigation of aiming magnitude in isolation. Then, to meaningfully aggregate the data of multiple participants, we realigned the chronology of their aiming data such that each individual’s first post-”Aha!” trial was at trial *t* = 0. We can see in Figure 3B that aiming on trial *t-1* was tightly grouped near zero; indeed, in experiment 1b, two-sample t-tests revealed no significant increase in aiming magnitude during the pre-”Aha!” trial versus baseline aiming for any rotation size (ps > 0.433). The same analysis on experiment 2b, Bond & Taylor, and Brudner et al. datasets, separately, all replicate this finding for all rotation sizes (ps > 0.12).

This ubiquitous perseveration in baseline behaviour was followed by an abrupt single-trial shift to good magnitude performance; aim magnitudes on the “Aha!” trial in all experiments (also including Bond & Taylor and Brudner et al.) were similar to the ideal magnitude (ps > 0.16). However, some participants intermittently return to their baseline strategy during the early post-”Aha!” phase (e.g., Figure 3E). This meant that the aiming magnitude on the following eight trials was significantly below the ideal in many rotation groups of several experiments (e.g., Figure 3C).

The “Aha!” moment, which typically occurs after 1-10 rotation trials (Figure 3D), is followed by a period not only of sometimes returning to one’s baseline strategy, but also of common sign-flip errors. A sign-flip error is where the magnitude of aiming appears to be a roughly correct noisy estimate, but the sign of the aim is in the wrong direction to compensate for the rotation. As can be seen in Figure 3F, these sign-flip errors are frequent shortly after the “Aha!” moment but get less frequent over time.

Taken together, these findings indicate that there is a clear discontinuity in aiming behaviour that occurs after some variable duration of baseline-like aiming. Aiming before this “Aha!” moment is not different from baseline; after the “Aha!” moment, the participants’ newly generated aiming strategy generally exhibits the correct magnitude of compensation (albeit in a noisy manner) but often requires additional learning to achieve a globally correct compensatory sign. Additionally, intermittent returns to the baseline strategy occur for some people in this post-”Aha!” phase.

### Is there any learning or exploration before the “Aha!” moment?

So far, the data strongly suggest the presence of an “Aha!” moment that manifests in an abrupt single-trial shift to a new, distinct strategy. However, it may be that learning or exploration do occur before the “Aha!” moment, but in a subtle manner. This might indicate that the true learning process is actually one of trial and error, rather than the result of an “Aha!” moment. To investigate this possibility, we now quantitatively test for any signs of learning or exploration before the “Aha!” moment.

We first looked at exploration. We took the pre-”Aha!” rotation-phase aiming data and tested whether there was any difference in trial-to-trial aiming variability compared to baseline aiming (Figure 4A). To avoid any sample-size bias effects, we ensured the same number of trials from each phase were considered; for example, if there were only four pre-”Aha!” rotation phase trials, then we also only considered the final four baseline trials for that participant. For this analysis we excluded participants with only one pre-”Aha!” rotation-phase trials (as no measure of variability can be calculated from a single trial). If exploration were present, we would expect an increase in aiming variability versus baseline. Pre-”Aha!” aiming variability was not different to baseline for any dataset (p’s > .075) apart from Brudner, where a significant *decrease* was observed (baseline: 8.39° [CI95%: 6.5°, 11.88°]; pre-”Aha!”: 5.14° [CI95%: 4.03°, 6.29°]; t(19) = 2.24, p = 0.038), likely because two participants showed abnormally high variability during baseline (Figure 4A) or because participants continued to improve their baseline-strategy execution during the pre-”Aha!” rotation phase.

**Figure 4.**
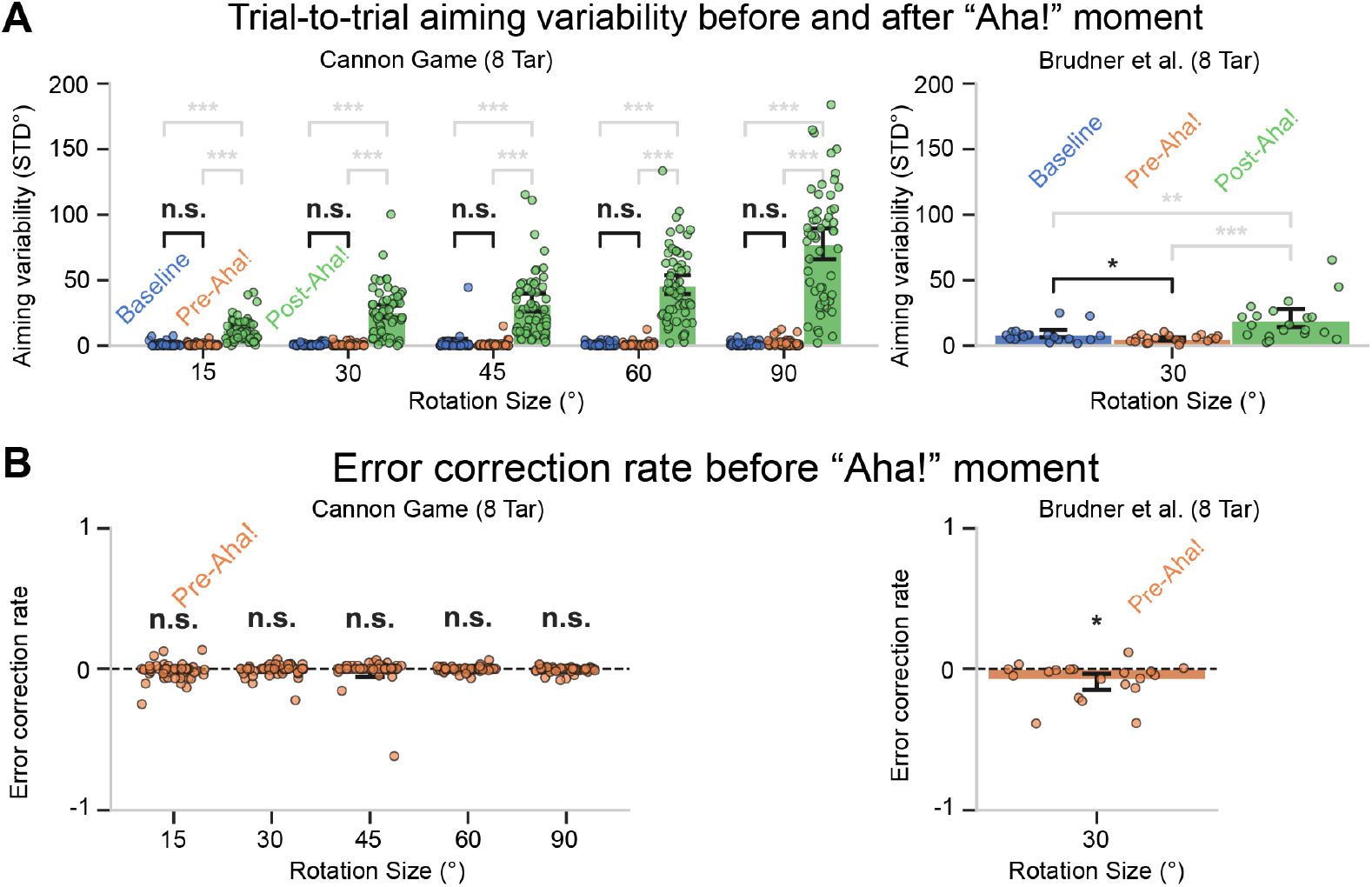
Pre-”Aha!” rotation-phase aiming behaviour does not differ from baseline aiming behaviour. (A) Trial-to-trial aiming variability during baseline (blue), pre-”Aha!” rotation-phase (orange), and post-”Aha!” rotation-phase (green) for vanilla Cannon Game (left) and Brudner et al (right). Significance results of paired t-tests are shown; the key test result of baseline versus pre-”Aha!” rotation-phase aiming variability is in black. (B) Error correction rate during pre-”Aha!” rotation-phase for vanilla Cannon Game (left) and Brudner et al (right). Significance results of one-sample t-tests are shown. These analyses used matched-window sizes, such that if the participant has n pre-”Aha!” rotation-phase trials, then only the last n baseline and first n post-”Aha!” trials are used. Only participants with at least two pre-”Aha!” rotation trials were included in the variability and error correction analyses,

We also submitted these pre-”Aha!” rotation-phase data to a test of significant error correction rate (i.e., the average trial-wise correction for errors during the pre-”Aha!” phase) using one-sample t-tests against zero (Figure 4B). If any learning were present, we would expect a positive error correction rate. The pre-”Aha!” correction rate was not different than 0 in any rotation size of any dataset (ps > 0.09), except Brudner et al. which was significantly *negative* (−0.081, CI95%: −0.15, −0.05, t(19) = 2.79, p = 0.012)), indicating a small increase in error during the pre-”Aha!” phase.

Together, these results reinforced our earlier findings this pre-”Aha!” rotation-phase aiming appears to be no different from baseline aiming and exhibits no signs of exploration or error correction.

### Model-based analysis

Having established the presence of an “Aha!” moment which is preceded by baseline behaviour and succeeded by a noisy, sometimes only partially correct strategy, we next employed model-based analyses to more directly test our “Aha!” hypothesis against traditional explanations of strategic behaviour in VMR tasks. We used three models in this analysis: to model gradual, error-based learning, we used a single-process SSM (which yields a better BIC than a dual-process SSM; Figure S3); for trial-and-error learning we propose a novel variant of the Q-learning algorithm which learns separate Q-functions for sign and magnitude, and carries out off-policy updates to overcome the sample inefficiency common in motor learning RL models; and to model the “Aha!” moment, we embed our novel modified Q-learning algorithm within a two-state HMM. This “Aha!” model must infer that the prediction error is relevant before it learns to compensate. This process typically results in a learning explosion, over a single trial, after some arbitrary period of baseline perseverance (e.g., Figure S1; also see Methods section for model algorithms).

We fit all three models to the first block of each individual’s aiming data in all experiments, as well as to the Bond & Taylor dataset and the Brudner et al. dataset (see methods for details). Parameter fits for all models can be seen in Figure 45E. Of the 777 participants across all datasets, 752 participants were best fit by the “Aha!” model according to BIC scores (we also fit the models to Ding et al. (2025), a recent study isolating strategic behaviour through delayed feedback, and find the “Aha!” model yielded the lowest BIC for 549 out of 560 participants; Figure S4G.) As can be seen in Figure 5A, the “Aha!” model replicates the abrupt shift from baseline aiming to correctly adjusting its compensatory magnitude for the noisily estimated feedback rotation (also see Figures S4A & B). The “Aha!” model also reproduces the sign uncertainty and intermittent recurrence of the baseline strategy, while the SSM fails to display any of these behaviours (Figure 5B & C).

**Figure 5.**
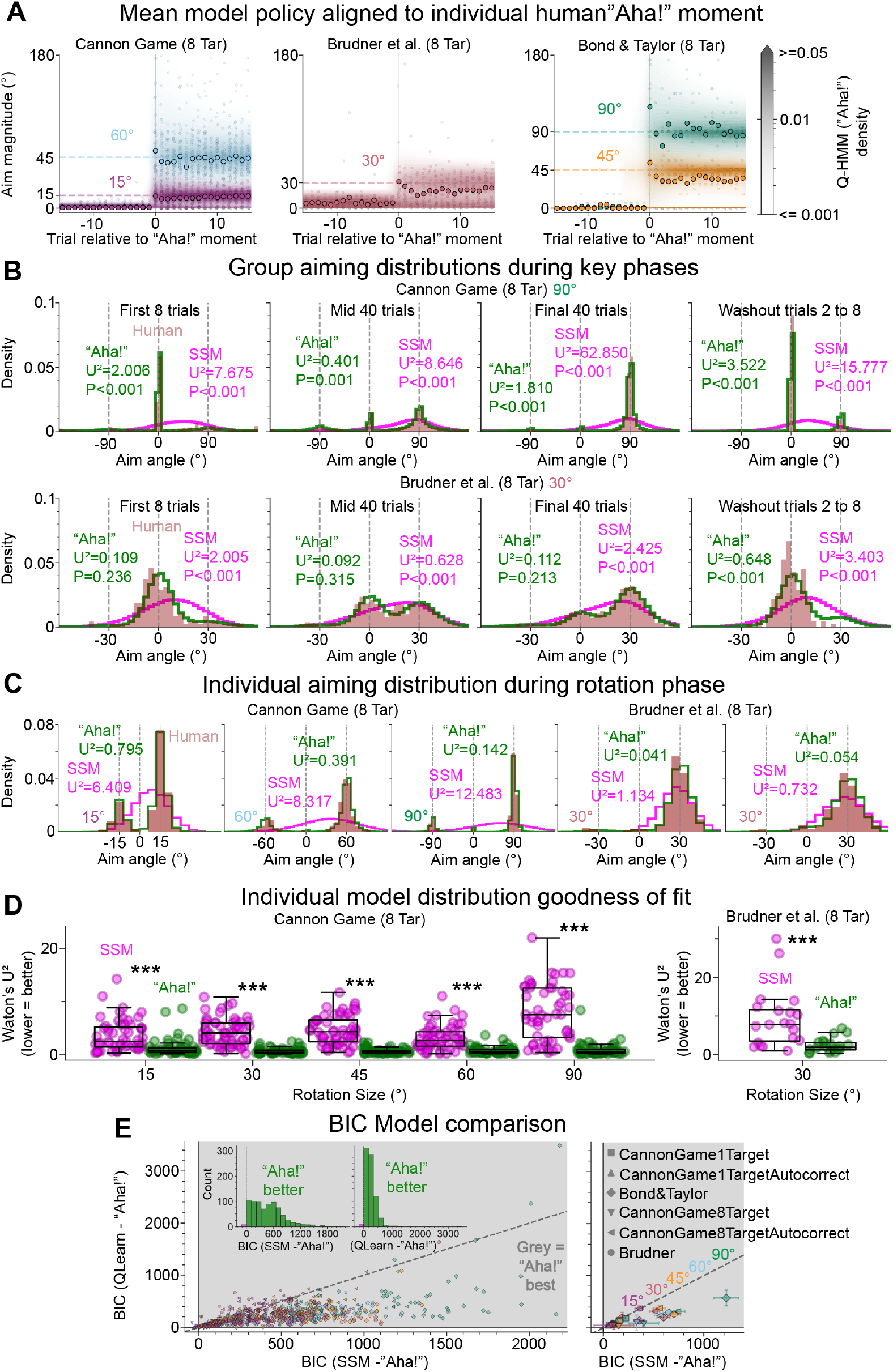
Human aiming behaviour is more like the “Aha!” model than to gradual error-based learning or Q-learning. (A) Human aiming magnitude (points) and mean “Aha!” model predictive policy distribution (contours) over time, for vanilla Cannon Game (left), Brudner et al (middle), and Bond & Taylor (2015; right), using data realigned such that trial 0 is the first trial after each individual human’s algorithmically detected “Aha!” moment. (B) Group-level distributions of signed human aiming (red), “Aha!” model predictive density (green), and SSM predictive density (magenta), during the first eight rotation trials (left), the next 40 trials (middle left), the final 40 rotation trials (middle right), and the second to eighth washout trials (right), for vanilla Cannon Game (top) and Brudner et al (bottom). Watson’s U2 test statistic is printed on each panel, for each model versus the human aims, which measures deviance from homogeneity between two circular distributions (lower = better). (C) Individual human signed aiming distributions (red), “Aha!” model predictive density (green), and SSM predictive density (magenta), for the entire rotation phase, for vanilla Cannon Game (left) and Brudner et al (right). (D) Watson’s U2 test statistic values for all individuals in vanilla Cannon Game (left) and Brudner et al (right), for the “Aha!” model (green) and the SSM (magenta). Significance results for paired t-tests between SSM and “Aha!” models’ U2 are shown (black stars). (E) Relative BIC scores for the three fitted models for individuals (left) and group means (right), for all six datasets (vanilla Cannon Game 1-target (square) & 8-target (diamond); autocorrecting Cannon Game 1-target (up-triangle) & 8-target (left-triangle); Brudner et al (circle); Bond & Taylor (down-triangle). The grey quadrant denotes a better fit for the “Aha!” model than for the SSM or the Q-learning model. The identity line indicates the threshold for Q-learning versus SSM (below the identity line denotes that Q-learning fit better than the SSM). Inset: number of participants within each range of relative BIC scores for the SSM versus “Aha!” (left) and Q-learning versus “Aha!” (right); green = “Aha!” model better, and magenta = “Aha!” model worse. Error bars are 95% confidence intervals.

To quantify the fidelity of each model to the human data, we submit the mean rotation-phase predictive policy of each model to a Watson’s U^2^ test against the respective human distribution of rotation-phase aims. This is a non-parametric test of two independent samples of data coming from a common circular distribution. This test is very sensitive to subtle differences, so a lack of significant difference for most participants was beyond the scope of this endeavour. However, the U^2^ metric is still a useful measure of similarity between two cumulative distribution functions (CDF), and it naturally handles circular data. Watson’s U^2^ integrates the squared deviations between both CDFs; lower is better. The Bond & Taylor dataset was excluded from this distributional analysis as the possible aim location reports were in increments of 5.625°, which causes a very specific and unnatural distribution that is very different from any model or other dataset.

We found that Watson’s U^2^ favoured the “Aha!” model across all rotation sizes in experiment 1b (ts > 7.34, ps < 0.001, Cohen’s ds > 0.99), experiment 2b (ts > 5.94, ps < 0.001, Cohen’s ds > 0.78), Brudner et al (t = 4.03, p < 0.001, Cohen’s d = 0.9), and Ding et al (ts = 15.4, p < 0.001, Cohen’s ds > 0.65).

These findings, combined with the visualisations in Figures 5 & S4, further evidence the presence of an “Aha!” moment. More specifically, they indicate that our “Aha!” model accurately captures much of human behaviour, whereas gradual error-minimisation or mere trial-and-error learning do not.

## Discussion

Strategic behaviour in VMR tasks is commonly characterised as either a gradual error-minimisation process, or as a process of trial-and-error learning. We sought to test these hypotheses on isolated strategic behaviour. We found that individual learning functions often do not gradually increase towards the ideal aim angle, nor do they typically appear to explore various strategies as predicted by typical trial-and-error learning accounts. Instead, it appears that most individuals typically perseverate in aiming at the target before experiencing an “Aha!” moment and suddenly shifting their aiming magnitude to compensate for a noisily estimated feedback perturbation. The correct sign often then takes more time to be learned after the “Aha!” moment. Some people do adopt a strategy which is entirely unlike the optimal strategy, but they still exhibit the same “Aha!” moment before its adoption.

We re-examined data from a delayed-feedback VMR task, which had no aim reports or prompts (Brudner et al., 2016), and uncovered the same hallmarks of an “Aha!” moment. We find further support for the “Aha!” hypothesis in a recent dataset by Ding et al. (2025), though the “Aha!” moment was expedited (and likely corrupted) by an explicit instruction to explore aiming strategies. Model-based analyses corroborate the presence of an “Aha!” moment.

### Strategic behaviour without motor contamination

We isolated strategic behaviour in a cannon-shooting game, removed from motor contaminants. In experiments 1a and 1b, with one target and eight targets respectively, we established that behaviour was generally consistent with that in VMR literature. We replicated the gradual, monotonic improvement in group-level performance followed by asymptote (Figure 1b).

To bring the aiming requirements of our task closer to those in a VMR task, we added an autocorrection module into the cannon. This covertly and gradually learned to reduce the discrepancy between the aim and feedback in a similar manner to implicit adaptation (Figure 2a & b; see Methods for details). This autocorrection used plan-based generalisation of implicit adaptation (Day et al., 2016; McDougle et al., 2017; Schween et al., 2018). We simulated the effects of using target-based generalisation instead and found no major differences (Figure S2).

We then characterised the degree of similarity between Cannon Game data and VMR data where performance is broken down into aim reports and implicit adaptation (Bond & Taylor, 2015). Aiming behaviour in experiment 2b did not differ from Bond & Taylor during the early, mid, or late rotation phases under any rotation size except 90°. This is consistent with strategic behaviour correcting for errors left by the sensorimotor system (Taylor & Ivry, 2011). We next defined three essential properties to characterise an individual’s learning curve, including: the cycle of peak aim; the mean aim during the peak cycle; the decrease in aim for the peak to final cycle. These properties were generally similar across experiments (e.g., Figure 2f).

Overall, behaviour appears to be generally similar across paradigms, prompting a deeper investigation into individual learning.

### The individual timecourse of aiming

At the group level, aiming increases gradually and monotonically towards the ideal aim location. The typical individual, however, tends to perseverate in aiming at the target until abruptly shifting their aim to approximate compensatory magnitude of the noisily estimated feedback rotation (e.g., Figure 3B). These disparate patterns of behaviour are partly reconciled by the fact that taking the mean over variably-delayed step-functions yields a smooth curve (Bahrick et al., 1957; Gallistel et al., 2004). Another contributing factor to this artefactual group-level curve is that participant’s typically take additional time after their “Aha!” moment to reliably use the correct sign, and they intermittently revert to baseline behaviour (Figure 3E & F).

### Alternative accounts of strategic behaviour

Strategic behaviour has traditionally been modelled as a gradual error-minimisation process via a SSM (Coltman et al., 2019; McDougle et al., 2015; Taylor & Ivry, 2011; Takiyama et al., 2015; Zhang et al., 2025). As can be seen in Figures 3A and S4A, there is a clear discontinuity in aiming strategy, and no sign of prior gradual error-minimisation. Viewing the data with our “Aha!”-aligned trial indices, we see that the same general absence of prior gradual learning at the group level (Figure 3C & S4B). While there may still be error-based fine-tuning of the strategy, strategy adoption seems discrete.

Another competing account is that strategic re-aiming in VMR tasks reflects a process of exploration until an efficacious action is found and can be exploited (e.g., Cashaback et al., 2017). This RL account predicts exploration of competing strategies before settling on the correct one. However, as can be seen in Figure 4 and Figure S4C & D, there is no evidence of an increase in trial-to-trial aiming variability, or error correction rate, during the pre-”Aha!” rotation phase. As such, we believe that the initial strategy adoption is the result of a changed internal representation (marked by the “Aha!” moment).

Finally, a recent account proposes that strategic behaviour reflects a process of hypothesis testing (Tsay et al., 2024; Ding et al., 2025). This entails the generation of hypotheses about the nature of the environment, and the testing of these hypotheses to gather information about their efficacy. As with the RL account, we do not view this as being at odds with our “Aha!” moment hypothesis; rather, they are focussed on different aspects of learning.

Central to our hypothesis is that the feedback error is considered uninformative to the agent’s goals until some moment of insight occurs and the participant’s internal representation of the task changes. This appears to typically manifest in a strategic solution which is correct, in terms of magnitude, for a noisily estimated perturbation (Figure 3B & 3C). The hypothesis-testing account is focussed on what happens after the “Aha!” moment. It may, of course, be the case that there is some covert process of hypothesis testing, not apparent in behaviour, that occurs prior to the “Aha!” moment. In any case, we do not see that as necessarily being contra to our account.

Some participants do initially adopt an incorrect compensatory magnitude. For example, in the 90° condition of experiment 1b, there is a small group of people who are clearly aiming ∼180° away from the target (Figure 3B). No other group of any experiment showed this phenomenon. This is still consistent with our “Aha!” moment hypothesis, though the “Aha!” moment led to the wrong strategy; in this case the participant must then change their strategy, which naturally invokes the RL or hypothesis-testing accounts. The sign learning may also reflect a process of hypothesis testing (or perhaps another “Aha!” moment). In this way, we see our account as focussing on initial strategy discovery, while the hypothesis-testing account is focussed more on what may happen after.

A much higher proportion (though still a minority) of participants from Ding et al. (2025) appear to adopt an incorrect strategy after their “Aha!” moment (e.g., aiming 180° away; Figure S4F. However, these participants were explicitly instructed to explore new aiming strategies to compensate for a change in action-feedback mapping during the rotation phase. This instruction appears to have expedited and corrupted the “Aha!” moment by encouraging premature strategy exploration (for example, 145 participants changed their strategy on the first rotation trial before observing any rotated feedback; Figure S4E).

Overall, our analyses indicate that the “Aha!” moment account explains much of the strategic behaviour in VMR tasks, leaving a relatively smaller portion for RL or hypothesis testing.

### The generative “Aha!” model

We developed a generative model that combines hierarchical off-policy Q-learning with a two-state HMM: in one state, the prediction error is considered irrelevant to the agent’s goals, and in the other state, the prediction error is considered relevant. In this way, the model can perseverate in baseline behaviour due to a strong prior that the prediction error is irrelevant. The dense rewards and off-policy nature of learning allow the model to shift from baseline aiming on trial *t* to the ideal aim on trial *t+1* (Figure S1). The off-policy updates allow the model to counterfactually generalise prediction error across the workspace, akin to counterfactual reasoning observed in humans (Byrne, 2002; Van Hoeck et al., 2015). It learns separate Q-functions for sign and magnitude, recapitulating the pattern of sign-flip errors in the post-”Aha!” rotation phase (Figures 5B & C and S4 H & I).

State inference within the model is subject to uncertainty over the prediction error. This context uncertainty increases with rotation size (Figure S6A), which aligns well with the fact that rotation size had no effect on the delay before an “Aha!” moment, despite the larger rotations providing a stronger error signal (Figure 3D). We know from Zhang et al. (2024) that the standard deviation of visual uncertainty increases in a roughly linear fashion (also see Klein & Levi., 1987; Levi et al., 1987). Indeed, we fit a linear function to context uncertainty and found a slope similar to that found by Zhang et al. (Figure S5).

We also see an exponential decay in inverse temperature as rotation size increases (Figure S6A). In RL, the inverse temperature defines the trade-off between exploration and exploitation. For our model, this means that larger rotation sizes, which elicit greater context uncertainty, see greater behavioural variability. However, it may be the case that the increased variability reflects the difficulty in accurately estimating the prediction error, or reflects bet-hedging against the uncertainty therein, rather than exploration over varied strategies. When present reward can augment future reward, hedging against uncertainty maximises the long-term reward in a varying environment (Philippi & Seger, 1989; Starrfelt & Kokko, 2012). For example, it is an optimal strategy for maximising long-term returns under variable financial market conditions (Markowitz, 1952; Jorion, 2009) or for maximising population fitness in dynamic adverse environments (Hopper, 1999; Olofsson et al., 2009).

Many natural human tasks do involve rewards which can augment future rewards. For example, any task that results in food, water, or social status thus provides sustenance to better perform subsequent tasks. Thus, it may be that the increase in exploration associated with an increase in context uncertainty reflects a form of hedging to maximise lifelong geometric-mean reward (also see Kaufmann et al. 2012 for proof of Thompson sampling optimality for cumulative regret minimisation). Work by Feng et al. (2021) is consistent with this hypothesis and finds that exploration is driven by the signal-to-noise ratio of reward information. Thus, we do not see good reason to believe that the variable compensatory magnitude reflects exploration of strategies.

The “Aha!” model isn’t intended to be a complete mechanistic account of the “Aha!” moment. Nevertheless, the model can generally capture the relevant phenomena covered in this work. While RL is essential to this model, the RL component relies on counterfactual reward to direct its behaviour, so we do not consider the initial strategy generation as being part of a trial-and-error learning process.

Out of 1337 participants in our modelling analysis, including data collected in VMR reaching tasks with no explicit aim reports, 1301 were best fit by our “Aha!” model (Figures 5E & S4G). Furthermore, only the “Aha!” model was able to reproduce the baseline perseveration along with the multimodal distribution of aims (e.g., Figures 5B & C; S4H & I). Taken together, we see this as overwhelming support for our “Aha!” model being superior to extant alternatives.

### The “Aha!” moment in the motor context

It is possible that our task yields the same behavioural patterns as in standard VMR tasks, but through distinct strategic processes. To investigate this concern, we carried out the same model-fitting analysis on pre-existing reaching data from a task without aim reports, and without any prompt that an aiming solution might exist or that one should consider aiming (Brudner et al., 2016). Instead, the task presented feedback after a delay period of 5 seconds, which is thought to inhibit implicit adaptation and emphasise strategic contributions (Schween & Hegele, 2017). All 20 participants were best fit by the “Aha!” model, and behaviour exhibited the same perseveration and abrupt shift (Figure 3C). Further, we see no reason, *a priori*, to think that strategic processing would be qualitatively—not just quantitatively—affected by this delay.

We also fit the models to recent data from Ding et al. (2025), where participants reached as in typical VMR tasks, though with explicit instruction to explore new strategies. The “Aha!” model still better explained the behaviour of 549 out of 560 participants, and it still matched the general behavioural patterns and distributions (Figure S4).

By incorporating our insights into strategic behaviour back into the motor adaptation context, we can attack the problem of implicit learning in new ways. For example, by using our generative “Aha!” model (or a successor) to explain away the portion of behaviour attributable to the strategic system, we may be able to investigate implicit adaptation in more ecologically relevant contexts than using error clamps or delayed feedback.

## Conclusion

Together, these findings indicate that strategic behaviour, both in the Cannon Game and in VMR reaching tasks, is not a gradual error-minimisation process or mere trial-and-error learning. Instead, there is perseveration of baseline behaviour, followed by an “Aha!” moment which marks a dramatic single-trial shift in behaviour. This typically manifests in a correct compensatory magnitude for a noisily estimated rotation but reliably correct sign behaviour takes longer to learn. This “Aha!” moment appears to eat much of the lunch of the trial-and-error accounts. This is important not only for understanding how strategies are acquired and evolve over time, but also in our attempt to understand implicit adaptation and motor behaviour more generally.

Future work should try to better understand the processes that lead to this sudden shift in behaviour. Is there a fundamental change in the representation of the task, or does the preceding perseveration reflect the time taken to infer the presence of a new context? Answering this would also expedite the motor rehabilitation process by helping patients to learn strategic behaviours more quickly and effectively.

## Acknowledgements

We thank Jordan Taylor for providing the raw data used to compare Gannon Game with VMR data (Bond & Taylor, 2015; Brudner et al., 2016).

## Conflict of interest

The authors declare no conflicts of interest.

## Financial support

M.T. was supported by an ESRC White Rose Doctoral Training Partnership Scholarship.

F.M. was supported by the UK Research and Innovation Biotechnology and Biological Sciences Research Council (BB/X008428/1) and the National Institute for Health and Care Research (NIHR) Leeds Biomedical Research Centre (NIHR203331).

## Data availability

Data and code will be made available upon acceptance on the Open Science Framework.

## Ethics statement

Ethical approval was obtained from the University of Leeds school of Psychology Research Ethics Committee (ref PSYC-81). All participants gave informed consent via a Qualtrics web form prior to starting the experiments.

## Author contributions

MT: Conceptualisation, Methodology, Software, Formal analysis, Investigation, Modelling, Visualisation, Writing - Original Draft, Writing - Review & Editing. MW: Validation, Writing - Review & Editing. CC: Supervision, Writing - Review & Editing. MMW: Supervision, Writing - Review & Editing, Funding acquisition. FM: Methodology, Supervision, Writing - Review & Editing, Funding acquisition. JRM: Conceptualisation, Methodology, Supervision, Writing - Review & Editing, Funding acquisition.

## Supplementary material

**Figure S1.**
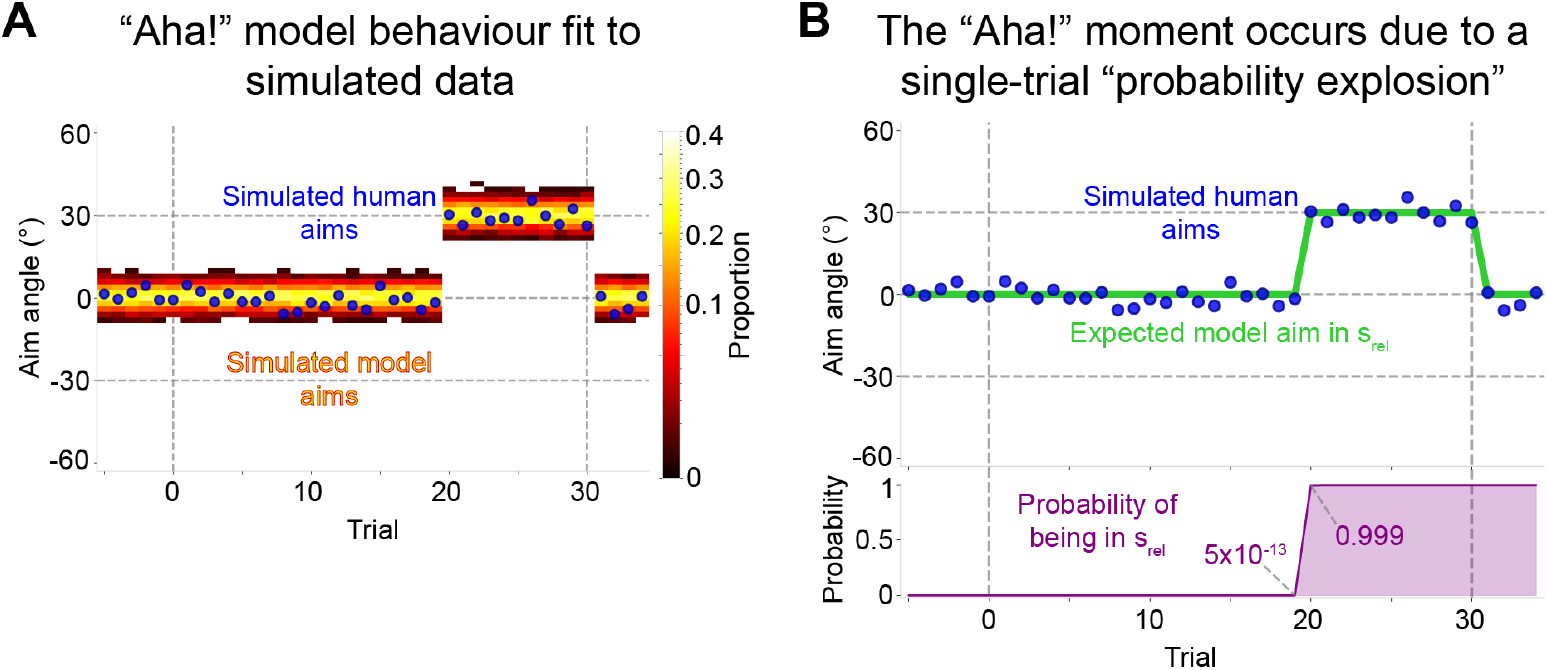
The “Aha!” model produces an “Aha!” moment-like jump to good performance through a single-trial “probability explosion”, where the probability of the prediction error being relevant jumps from effectively zero to effectively one. (A) Timecourse of simulated human aims (blue) with simulated model aims (‘hot’ heatmap; 500 draws per trial). (B) Top: Timecourse of simulated human aims (blue) with expected model aim in *s*_*rel*_ (i.e., 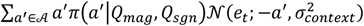; green). The model doesn’t behave according to a point estimate, but we use it here for illustrative purposes. Bottom: The probability that the prediction error is relevant to the agent’s goals (i.e., **P**_***t***_(*S*_*t*_ = *s*_*rel*_); purple). Vertical dashed lines indicate the first trial of the rotation phase and the first trial of the washout phase.

**Figure S2.**
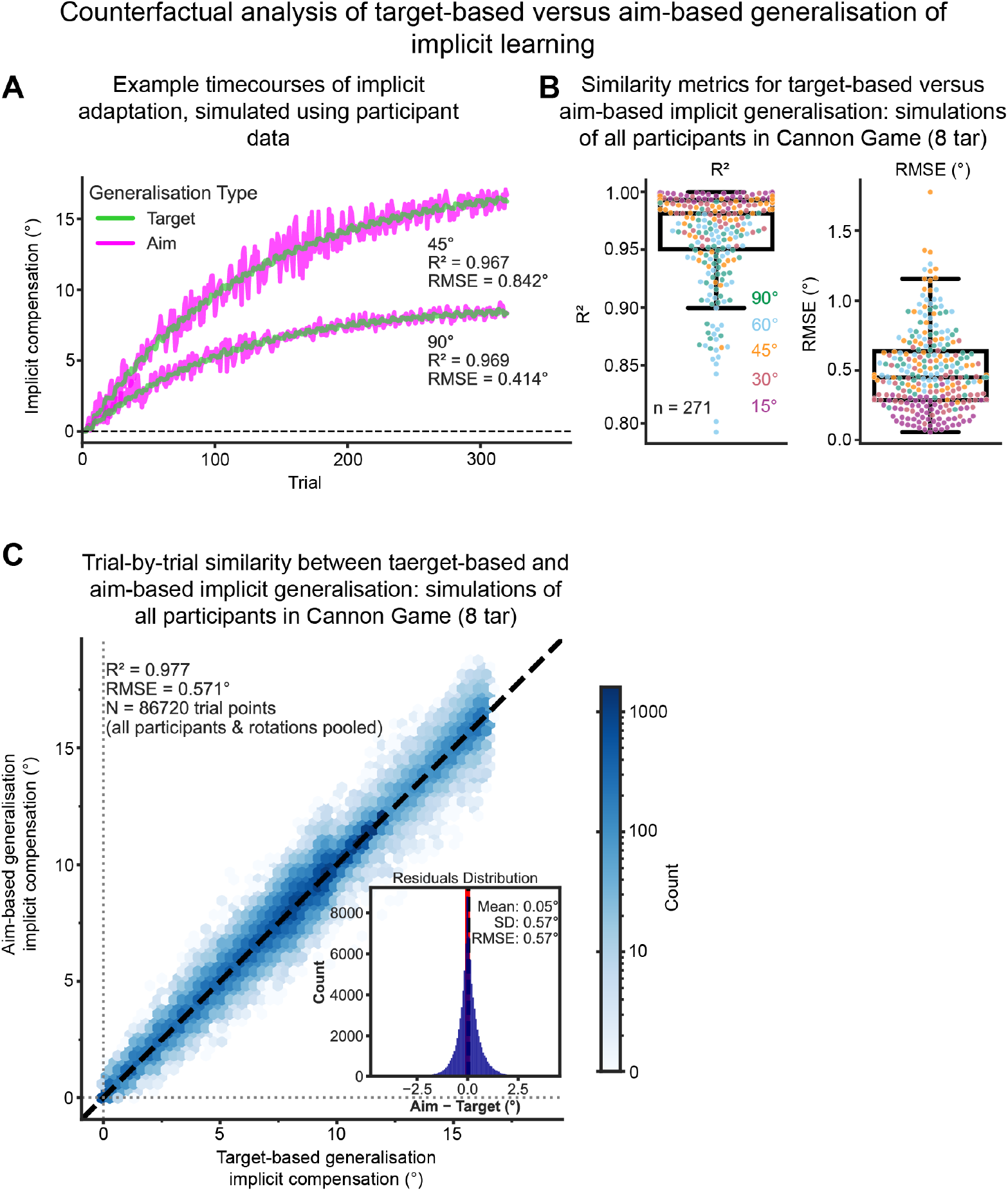
Target-based generalisation of implicit learning is similar to aim-based generalisation of implicit learning. (A) Example timecourses from vanilla Cannon Game (8-target) for actually-observed aim-based generalisation of implicit learning (green) and simulated target-based generalisation (magenta). (B) Per-participant R^2 (left) and RMSE (right), between their actually-observed aim-based generalising implicit learning and the simulated target-based generalising implicit learning, for all participants in the 8-target vanilla Cannon Game. (C) Trial-wise discrepancy between aim-based generalising implicit learning and target-based generalising implicit learning, pooling together all rotation trials from all participants in all rotation groups of vanilla Cannon Game. Identity line = both types of implicit learning are identical. (Note the log scale.) Inset: residuals over all rotation trials.

**Figure S3.**
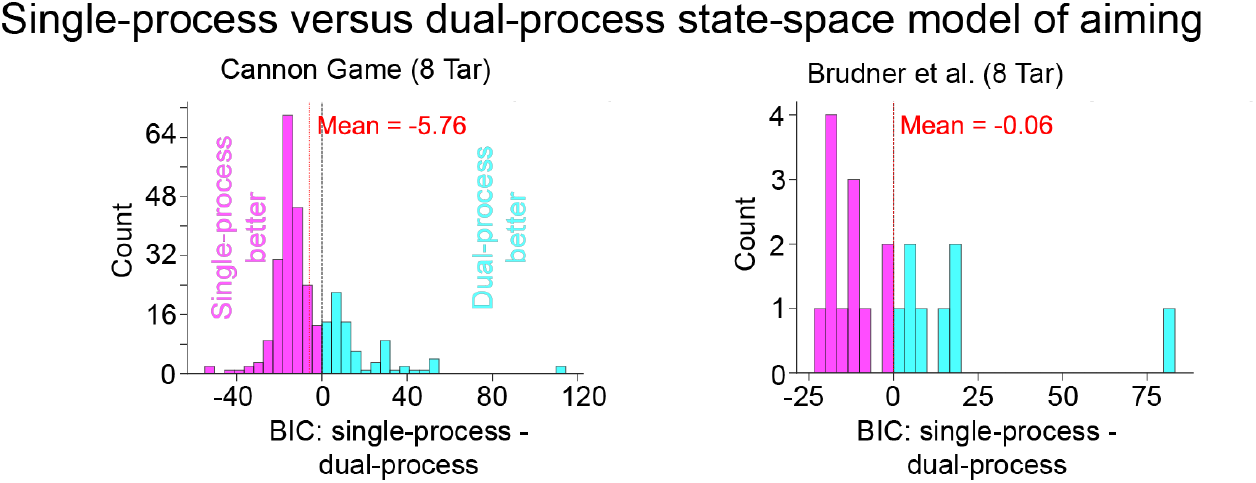
A dual-process state-space model offers no better fit to aiming behaviour, on average, than a single-process state-space model. Distribution of per-participant relative BIC values, for single-process versus dual-process SSM, for vanilla Cannon Game (left) and Brudner et al (right). Negative (magenta) denotes a better fit for the single-process SSM, while positive (cyan) denotes a better fit for the dual-process SSM.

**Figure S4.**
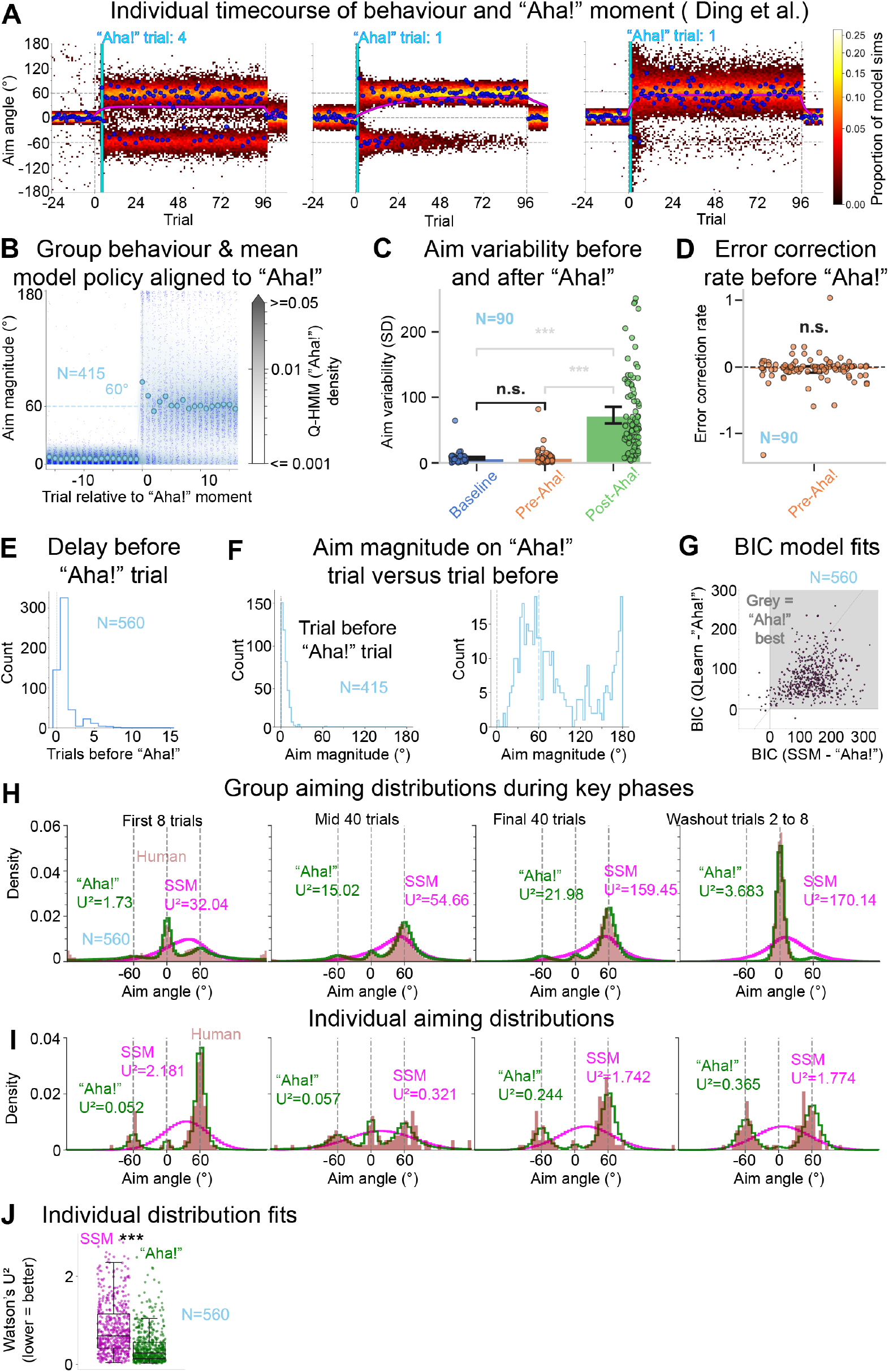
Participants in Ding et al (2025) also appear to behave consistently with the “Aha!” moment hypothesis. (A) Individual timecourses of behaviour for three participants; human aiming behaviour (blue), “Aha!” model samples (500 per trial) in ‘hot’ heatmap, SSM mean aim (solid magenta line), and the algorithmically detected “Aha!” trial denoted by the vertical cyan line. (B) Human group behaviour (points) with mean “Aha!” model policy (contours) for aim magnitude, with each indivdual’s behaviour realigned to their respective “Aha!” trial being set to zero. Participants who exhibited a changepoint on the first rotation trial (i.e., before having observed rotated feedback) were excluded from this analysis. (C) Trial-to-trial aiming variability during baseline (blue), pre-”Aha!” rotation-phase (orange), and post-”Aha!” rotation-phase (green). Significance results of paired t-tests are shown; the key test result of baseline versus pre-”Aha!” rotation-phase aiming variability is in black. (D) Error correction rate during pre-”Aha!” rotation phase. Significance results of one-sample t-tests are shown. These analyses used matched-window sizes, such that if the participant has n pre-”Aha!” rotation-phase trials, then only the last n baseline and first n post-”Aha!” trials are used. Only participants with at least two pre-”Aha!” rotation trials were included in the variability and error correction analyses, (E) The humber of rotation trials before each individual’s “Aha!” moment. (F) Aim magnitudes for the trial immediately preceding their respective “Aha!” moment (left) and on the subsequent trial (right). Participants who exhibited a changepoint on the first rotation trial (i.e., before having observed rotated feedback) were excluded from this analysis. (G) Relative BIC model fits. The grey quadrant denotes a better fit for the “Aha!” model than for the SSM or the Q-learning model. The identity line indicates the threshold for Q-learning versus SSM (below the identity line denotes that Q-learning fits better than the SSM). (H) Group-level distributions of signed human aiming (red), “Aha!” model predictive density (green), and SSM predictive density (magenta), during the first eight rotation trials (left), the next 40 trials (middle left), the final 40 rotation trials (middle right), and the second to eighth washout trials (right). Watson’s U2 test statistic is printed on each panel, for each model versus the human aims, which measures deviance from homogeneity between two circular distributions (lower = better). (I) Individual human signed aiming distributions (red), “Aha!” model predictive density (green), and SSM predictive density (magenta), for the entire rotation phase. (J) Watson’s U2 test statistic values of all individuals for the “Aha!” model (green) and the SSM (magenta). Significance results for paired t-tests between SSM and “Aha!” models’ U2 are shown (black stars).

**Figure S5.**
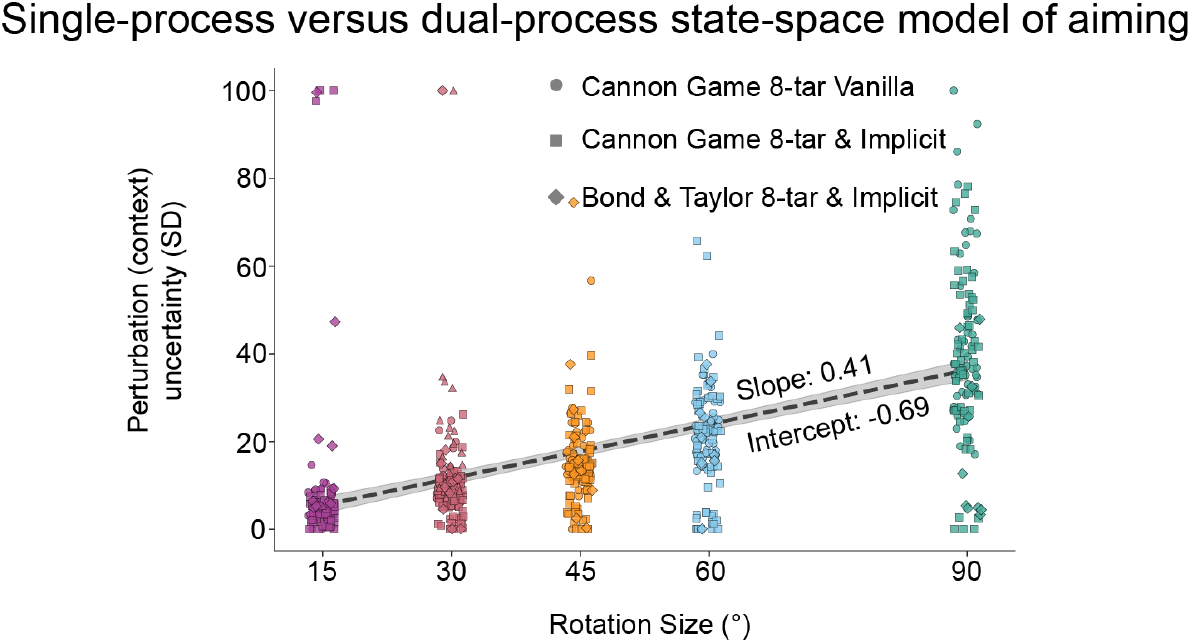
Linear fit to uncertainty in perturbation (context in “Aha!” model) by rotation size. All data from the 8-target Cannon Game experiments, as well as from Bond & Taylor (2015), are pooled together for this analysis. Error bands = 95% confidence intervals.

**Figure S6.**
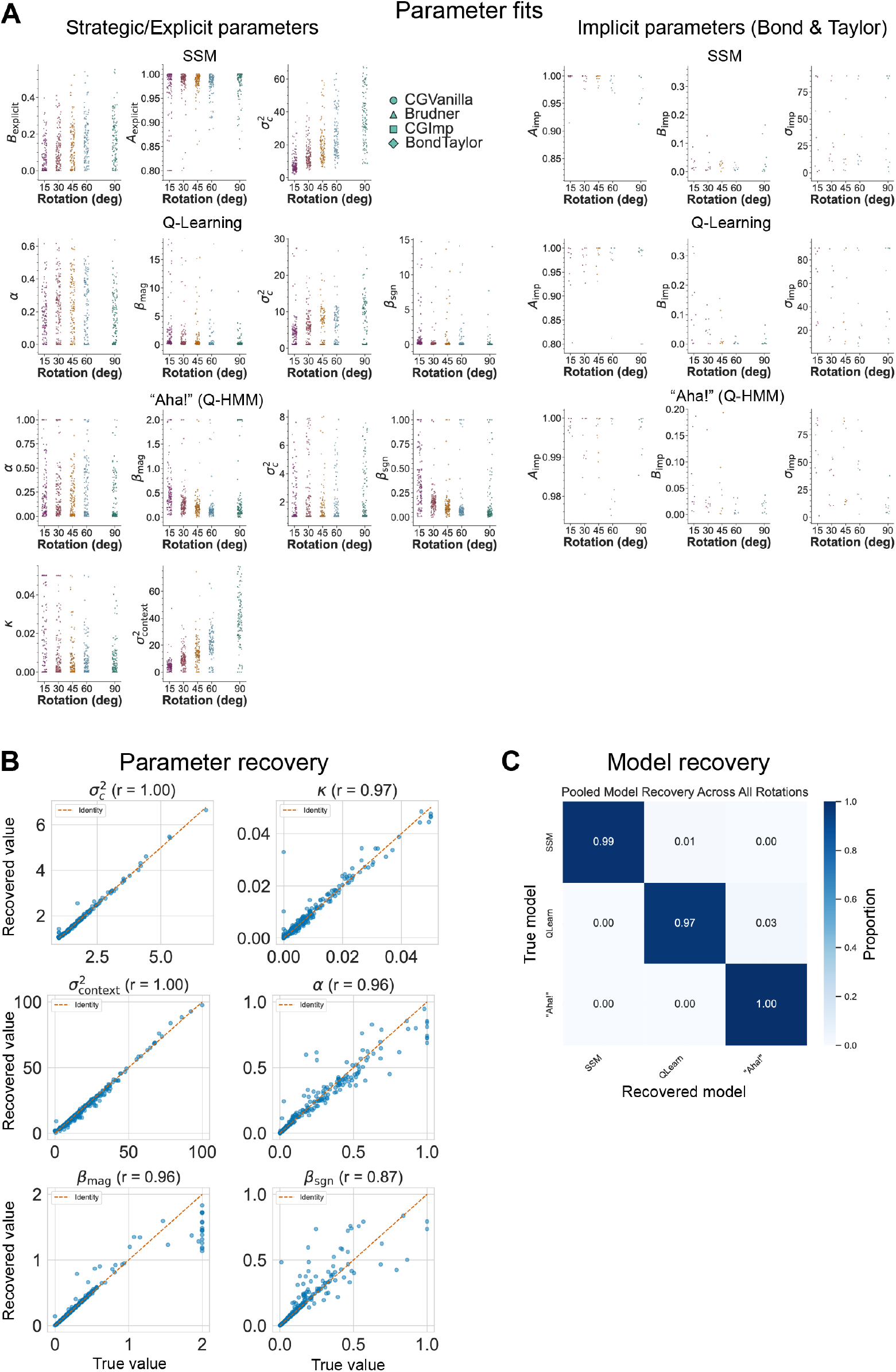
(A) Aiming model parameter fits (left), and implicit state-space model parameter fits (right). (B) Parameter recovery for the “Aha!” model. We used the parameters of all human participants from each rotation group of the 8-target vanilla Cannon Game dataset; we used each set of parameters to simulate twelve separate runs. These 3372 simulated participants were then refit using the “Aha!” model 200 times each. We used the mean parameter fits over the 12 runs for each participant. (C) Model recovery for each model. We used the parameter sets of every participant in the 8-target vanilla Cannon Game dataset to each simulate one run with each model. In this way, every model was then refit to the 843 simulated participants 250 times each.

## Notes

### Competing Interest Statement

The authors have declared no competing interest.

### Summary of Updates

Substantial revision of the paper, including a new generative model, new figures 3-5, revised discussion.

## References

Allen, K. R., Smith, K. A., & Tenenbaum, J. B. (2020). Rapid trial-and-error learning with simulation supports flexible tool use and physical reasoning. Proceedings of the National Academy of Sciences, 117(47), 29302–29310. 10.1073/pnas.1912341117

Bahrick, H. P., Fitts, P. M., & Briggs, G. E. (1957). Learning curves—Facts or artifacts? Psychological Bulletin, 54(3), 256–268. 10.1037/h0040313

Bond, K. M., & Taylor, J. A. (2015). Flexible explicit but rigid implicit learning in a visuomotor adaptation task. Journal of Neurophysiology, 113(10), 3836–3849. 10.1152/jn.00009.2015

Bowden, E. M., & Jung-Beeman, M. (2003). Aha! Insight experience correlates with solution activation in the right hemisphere. Psychonomic Bulletin & Review, 10(3), 730–737. 10.3758/BF03196539

Brudner, S. N., Kethidi, N., Graeupner, D., Ivry, R. B., & Taylor, J. A. (2016). Delayed feedback during sensorimotor learning selectively disrupts adaptation but not strategy use. Journal of Neurophysiology, 115(3), 1499–1511. 10.1152/jn.00066.2015

Byrne, R. M. (2002). Mental models and counterfactual thoughts about what might have been. Trends in cognitive sciences, 6(10), 426–431. 10.1016/S1364-6613(02)01974-5

Carpenter, W. (2020). The Aha! Moment: The Science Behind Creative Insights. In S. Manuel Brito (Ed.), Toward Super-Creativity—Improving Creativity in Humans, Machines, and Human—Machine Collaborations. IntechOpen. 10.5772/intechopen.84973

Cashaback, J. G. A., Lao, C. K., Palidis, D. J., Coltman, S. K., McGregor, H. R., & Gribble, P. L. (2019). The gradient of the reinforcement landscape influences sensorimotor learning. PLOS Computational Biology, 15(3), e1006839. 10.1371/journal.pcbi.1006839

Cohen, J. D., McClure, S. M., & Yu, A. J. (2007). Should I stay or should I go? How the human brain manages the trade-off between exploitation and exploration. Philosophical Transactions of the Royal Society B: Biological Sciences, 362(1481), 933–942. 10.1098/rstb.2007.2098

Collins, A., & Koechlin, E. (2012). Reasoning, Learning, and Creativity: Frontal Lobe Function and Human Decision-Making. PLoS Biology, 10(3), e1001293. 10.1371/journal.pbio.1001293

Coltman, S. K., Cashaback, J. G. A., & Gribble, P. L. (2019). Both fast and slow learning processes contribute to savings following sensorimotor adaptation. Journal of Neurophysiology, 121(4), 1575–1583. 10.1152/jn.00794.2018

Darshan, R., Leblois, A., & Hansel, D. (2014). Interference and Shaping in Sensorimotor Adaptations with Rewards. PLoS Computational Biology, 10(1), e1003377. 10.1371/journal.pcbi.1003377

Day, K. A., Roemmich, R. T., Taylor, J. A., & Bastian, A. J. (2016). Visuomotor learning generalizes around the intended movement. eneuro, 3(2). 10.1523/ENEURO.0005-16.2016

Ding, W., Niyogi, A., & Tsay, J. S. (2025). Hypothesis Testing Governs an Efficiency– Flexibility Trade-off in Strategic Motor Learning. bioRxiv, 2025–11

Donoso, M., Collins, A. G. E., & Koechlin, E. (2014). Foundations of human reasoning in the prefrontal cortex. Science, 344(6191), 1481–1486. 10.1126/science.1252254

Feng, S. F., Wang, S., Zarnescu, S., & Wilson, R. C. (2021). The dynamics of explore– exploit decisions reveal a signal-to-noise mechanism for random exploration. Scientific reports, 11(1), 3077. 10.1038/s41598-021-82530-8

Foster, D. J. (2017). Replay Comes of Age. Annual Review of Neuroscience, 40(1), 581–602. 10.1146/annurev-neuro-072116-031538

Gallistel, C. R., Fairhurst, S., & Balsam, P. (2004). The learning curve: Implications of a quantitative analysis. Proceedings of the National Academy of Sciences, 101(36), 13124–13131. 10.1073/pnas.0404965101

Griffiths, T. L., Callaway, F., Chang, M. B., Grant, E., Krueger, P. M., & Lieder, F. (2019). Doing more with less: Meta-reasoning and meta-learning in humans and machines. Current Opinion in Behavioral Sciences, 29, 24–30. 10.1016/j.cobeha.2019.01.005

Griffiths, T. L., Chater, N., Kemp, C., Perfors, A., & Tenenbaum, J. B. (2010). Probabilistic models of cognition: Exploring representations and inductive biases. Trends in Cognitive Sciences, 14(8), 357–364. 10.1016/j.tics.2010.05.004

Hegele, M., & Heuer, H. (2010). Implicit and explicit components of dual adaptation to visuomotor rotations. Consciousness and Cognition, 19(4), 906–917. 10.1016/j.concog.2010.05.005

Hopper, K. R. (1999). Risk-spreading and bet-hedging in insect population biology. Annual review of entomology, 44(1), 535–560. 10.1146/annurev.ento.44.1.535

Jones, G. (2003). Testing two cognitive theories of insight. Journal of Experimental Psychology: Learning, Memory, and Cognition, 29(5), 1017–1027. 10.1037/0278-7393.29.5.1017

Jorion, P. (2009). Risk management lessons from the credit crisis. European Financial Management, 15(5), 923–933. 10.1111/j.1468-036X.2009.00507.x

Jung-Beeman, M., Bowden, E. M., Haberman, J., Frymiare, J. L., Arambel-Liu, S., Greenblatt, R., Reber, P. J., & Kounios, J. (2004). Neural Activity When People Solve Verbal Problems with Insight. PLoS Biology, 2(4), e97. 10.1371/journal.pbio.0020097

Kaufmann, E., Korda, N., & Munos, R. (2012, October). Thompson sampling: An asymptotically optimal finite-time analysis. In International conference on algorithmic learning theory (pp. 199–213). Berlin, Heidelberg: Springer Berlin Heidelberg. 10.1007/978-3-642-34106-9_18

Killick, R., Fearnhead, P., & Eckley, I. A. (2012). Optimal Detection of Changepoints With a Linear Computational Cost. Journal of the American Statistical Association, 107(500), 1590–1598. 10.1080/01621459.2012.737745

Klein, S. A., & Levi, D. M. (1987). Position sense of the peripheral retina. Journal of the Optical Society of America A, 4(8), 1543–1553. 10.1364/JOSAA.4.001543

Knoblich, G., Ohlsson, S., Haider, H., & Rhenius, D. (1999). Constraint Relaxation and Chunk Decomposition in Insight Problem Solving. Journal of Experimental Psychology: Learning, Memory, and Cognition, 25(6), 1534–1555. 10.1037/0278-7393.25.6.1534

Knoblich, G., Ohlsson, S., & Raney, G. E. (2001). An eye movement study of insight problem solving. Memory & Cognition, 29(7), 1000–1009. 10.3758/BF03195762

Kounios, J., & Beeman, M. (2009). The Aha! Moment: The Cognitive Neuroscience of Insight. Current Directions in Psychological Science, 18(4), 210–216. 10.1111/j.1467-8721.2009.01638.x

Krakauer, J. W., Pine, Z. M., Ghilardi, M.-F., & Ghez, C. (2000). Learning of Visuomotor Transformations for Vectorial Planning of Reaching Trajectories. The Journal of Neuroscience, 20(23), 8916–8924. 10.1523/JNEUROSCI.20-23-08916.2000

Langsdorf, L., Maresch, J., Hegele, M., McDougle, S. D., & Schween, R. (2021). Prolonged response time helps eliminate residual errors in visuomotor adaptation. Psychonomic Bulletin & Review, 28(3), 834–844. 10.3758/s13423-020-01865-x

Levi, D. M., Klein, S. A., & Yap, Y. L. (1987). Positional uncertainty in peripheral and amblyopic vision. Vision research, 27(4), 581–597. 10.1016/0042-6989(87)90044-7

MacGregor, J. N., Ormerod, T. C., & Chronicle, E. P. (2001). Information processing and insight: A process model of performance on the nine-dot and related problems. Journal of Experimental Psychology: Learning, Memory, and Cognition, 27(1), 176–201. 10.1037/0278-7393.27.1.176

Maresch, J., Mudrik, L., & Donchin, O. (2021a). Measures of explicit and implicit in motor learning: What we know and what we don’t. Neuroscience & Biobehavioral Reviews, 128, 558–568. 10.1016/j.neubiorev.2021.06.037

Maresch, J., Werner, S., & Donchin, O. (2021b). Methods matter: Your measures of explicit and implicit processes in visuomotor adaptation affect your results. European Journal of Neuroscience, 53(2), 504–518. 10.1111/ejn.14945

Markowitz, H. M. (1952). Portfolio Selection, the journal of finance. 7 (1). N, 1, 71–91. 10.2307/2975974

Mattar, A. A. G., & Ostry, D. J. (2007). Modifiability of Generalization in Dynamics Learning. Journal of Neurophysiology, 98(6), 3321–3329. 10.1152/jn.00576.2007

McDougle, S. D., Bond, K. M., & Taylor, J. A. (2015). Explicit and Implicit Processes Constitute the Fast and Slow Processes of Sensorimotor Learning. Journal of Neuroscience, 35(26), 9568–9579. 10.1523/JNEUROSCI.5061-14.2015

McDougle, S. D., Bond, K. M., & Taylor, J. A. (2017). Implications of plan-based generalization in sensorimotor adaptation. Journal of Neurophysiology, 118(1), 383–393. 10.1152/jn.00974.2016

McDougle, S. D., Ivry, R. B., & Taylor, J. A. (2016). Taking Aim at the Cognitive Side of Learning in Sensorimotor Adaptation Tasks. Trends in Cognitive Sciences, 20(7), 535–544. 10.1016/j.tics.2016.05.002

McDougle, S. D., & Taylor, J. A. (2019). Dissociable cognitive strategies for sensorimotor learning. Nature communications, 10(1), 40. 10.1038/s41467-018-07941-0

Morehead, J. R., Butcher, P. A., & Taylor, J. A. (2011). Does Fast Learning Depend on Declarative Mechanisms? The Journal of Neuroscience, 31(14), 5184–5185. 10.1523/JNEUROSCI.0040-11.2011

Morehead, J. R., Qasim, S. E., Crossley, M. J., & Ivry, R. (2015). Savings upon Re-Aiming in Visuomotor Adaptation. Journal of Neuroscience, 35(42), 14386–14396. 10.1523/JNEUROSCI.1046-15.2015

Morehead, J. R., Taylor, J. A., Parvin, D. E., & Ivry, R. B. (2017). Characteristics of Implicit Sensorimotor Adaptation Revealed by Task-irrelevant Clamped Feedback. Journal of Cognitive Neuroscience, 29(6), 1061–1074. 10.1162/jocn_a_01108

Neville, K.-M., & Cressman, E. K. (2018). The influence of awareness on explicit and implicit contributions to visuomotor adaptation over time. Experimental Brain Research, 236(7), 2047–2059. 10.1007/s00221-018-5282-7

Novick, L. R., & Sherman, S. J. (2003). On the Nature of Insight Solutions: Evidence from Skill Differences in Anagram Solution. The Quarterly Journal of Experimental Psychology Section A, 56(2), 351–382. 10.1080/02724980244000288

Olofsson, H., Ripa, J., & Jonzén, N. (2009). Bet-hedging as an evolutionary game: the trade-off between egg size and number. Proceedings of the Royal Society B: Biological Sciences, 276(1669), 2963–2969. 10.1098/rspb.2009.0500

Philippi, T., & Seger, J. (1989). Hedging one’s evolutionary bets, revisited. Trends in ecology & evolution, 4(2), 41–44. 10.1016/0169-5347(89)90138-9

Piantadosi, S. T., Tenenbaum, J. B., & Goodman, N. D. (2016). The logical primitives of thought: Empirical foundations for compositional cognitive models. Psychological Review, 123(4), 392–424. 10.1037/a0039980

Salvi, C., Bricolo, E., Kounios, J., Bowden, E., & Beeman, M. (2016). Insight solutions are correct more often than analytic solutions. Thinking & Reasoning, 22(4), 443–460. 10.1080/13546783.2016.1141798

Sawyer, R. K. (2012). Explaining creativity: The science of human innovation, 2nd ed. (pp. xi, 555). Oxford University Press.

Schween, R., & Hegele, M. (2017). Feedback delay attenuates implicit but facilitates explicit adjustments to a visuomotor rotation. Neurobiology of Learning and Memory, 140, 124–133. 10.1016/j.nlm.2017.02.015

Schween, R., Taylor, J. A., & Hegele, M. (2018). Plan-based generalization shapes local implicit adaptation to opposing visuomotor transformations. Journal of neurophysiology, 120(6), 2775–2787. 10.1152/jn.00451.2018

Smith, R. W., & Kounios, J. (1996). Sudden insight: All-or-none processing revealed by speed–accuracy decomposition. Journal of Experimental Psychology: Learning, Memory, and Cognition, 22(6), 1443. 10.1037/0278-7393.22.6.1443

Sprugnoli, G., Rossi, S., Emmendorfer, A., Rossi, A., Liew, S.-L., Tatti, E., di Lorenzo, G., Pascual-Leone, A., & Santarnecchi, E. (2017). Neural correlates of Eureka moment. Intelligence, 62, 99–118. 10.1016/j.intell.2017.03.004

Starrfelt, J., & Kokko, H. (2012). Bet-hedging—a triple trade-off between means, variances and correlations. Biological Reviews, 87(3), 742–755. 10.1111/j.1469-185X.2012.00225.x

Sugiyama, T., Schweighofer, N., & Izawa, J. (2023). Reinforcement learning establishes a minimal metacognitive process to monitor and control motor learning performance. Nature communications, 14(1), 3988. 10.1038/s41467-023-39536-9

Takiyama, K., Hirashima, M., & Nozaki, D. (2015). Prospective errors determine motor learning. Nature communications, 6(1), 5925. 10.1038/ncomms6925

Taylor, J. A., & Ivry, R. B. (2011). Flexible Cognitive Strategies during Motor Learning. PLoS Computational Biology, 7(3), e1001096. 10.1371/journal.pcbi.1001096

Taylor, J. A., & Ivry, R. B. (2012). The role of strategies in motor learning: The role of strategies in motor learning. Annals of the New York Academy of Sciences, 1251(1), 1–12. 10.1111/j.1749-6632.2011.06430.x

Taylor, J. A., & Ivry, R. B. (2013). Context-dependent generalization. Frontiers in Human Neuroscience, 7. 10.3389/fnhum.2013.00171

Taylor, J. A., Klemfuss, N. M., & Ivry, R. B. (2010). An Explicit Strategy Prevails When the Cerebellum Fails to Compute Movement Errors. The Cerebellum, 9(4), 580–586. 10.1007/s12311-010-0201-x

Taylor, J. A., Krakauer, J. W., & Ivry, R. B. (2014). Explicit and Implicit Contributions to Learning in a Sensorimotor Adaptation Task. Journal of Neuroscience, 34(8), 3023–3032. 10.1523/JNEUROSCI.3619-13.2014

Trope, Y., & Liberman, N. (2010). Construal-level theory of psychological distance. Psychological Review, 117(2), 440–463. 10.1037/a0018963

Truong, C., Oudre, L., & Vayatis, N. (2020). Selective review of offline change point detection methods. Signal Processing, 167, 107299 10.1016/j.sigpro.2019.107299

Tsay, J. S., Kim, H. E., McDougle, S. D., Taylor, J. A., Haith, A., Avraham, G., Krakauer, J. W., Collins, A. G., & Ivry, R. B. (2024). Fundamental processes in sensorimotor learning: Reasoning, refinement, and retrieval. eLife, 13, e91839. 10.7554/eLife.91839

Van Hoeck, N., Watson, P. D., & Barbey, A. K. (2015). Cognitive neuroscience of human counterfactual reasoning. Frontiers in human neuroscience, 9, 420. 10.3389/fnhum.2015.00420

Vartanian, O., & Goel, V. (2007). Neural correlates of creative cognition. In C. Martindale, P. Locher, V. M. Petrov, & A. Berleant (Eds.), Evolutionary and neurocognitive approaches to aesthetics, creativity and the arts (pp. 195–207). Routledge.

Werner, S., Van Aken, B. C., Hulst, T., Frens, M. A., Van Der Geest, J. N., Strüder, H. K., & Donchin, O. (2015). Awareness of Sensorimotor Adaptation to Visual Rotations of Different Size. PLOS ONE, 10(4), e0123321. 10.1371/journal.pone.0123321

Wunderlich, K., Symmonds, M., Bossaerts, P., & Dolan, R. J. (2011). Hedging your bets by learning reward correlations in the human brain. Neuron, 71(6), 1141–1152. 10.1016/j.neuron.2011.07.025

Zhang, Z., Wang, H., Zhang, T., Nie, Z., & Wei, K. (2024). Perceptual error based on Bayesian cue combination drives implicit motor adaptation. Elife, 13, RP94608. 10.7554/eLife.94608

Zhang, X., Wu, W., & Wei, K. (2025). Perceptual Prediction Error Supports Implicit Process in Motor Learning. bioRxiv, 2025-04. 10.7554/eLife.94608.3

